# Integrating Mutation and Stop Signals for Improved RNA Structure Analysis and Insight Discovery

**DOI:** 10.1101/2025.10.08.681285

**Authors:** Tong Zhou, Ruobin Zhao, Shaozhen Yin, Yifan Hong, Junhao Wang, Xitong Liu, Qiuzhen Chen, Yuancheng Wang, Chang Liu, Lei Sun

## Abstract

**Background:** Understanding RNA structures is essential for exploring its diverse cellular roles. Chemical modification-based RNA structure probing remains a key approach to revealing RNA structures in complex *in vivo* conditions. Practically, chemical modifications generate both mutation and stop signals during reverse transcription within a single experiment. However, traditional analysis pipelines often rely on only one of the two signals without fully leveraging both.

**Results:** To address this, we developed a machine learning-based approach, STONE, that integrates both signals from a single experiment. STONE improved RNA structure analysis across multiple independent validation regions, notably 28S rRNA, viral RNA and regulatory RNAs. In genome-wide datasets, especially single-cell data, STONE significantly increased nucleotide coverage per transcript and improved gene detection reliability. In genome-wide datasets, especially single-cell data, STONE substantially enhanced structural information coverage at both transcript and nucleotide levels. This maximized signal utilization, yielding RNA structures in single-cell data comparable to those from bulk datasets. Furthermore, STONE-derived structural scores allow direct identification of RNA-binding protein binding sites directly from a single probing experiment, with results on RNAs such as U1 snRNA, RNase P RNA, and XIST lncRNA closely matching established techniques like CLIP-seq and RNP-MaP.

**Conclusions:** By integrating mutation and stop signals from one single experiment, STONE advances RNA structure analysis accuracy, extends nucleotide coverage, and facilitates complex RNA-protein interaction studies, broadening the method’s applications in RNA-based research.

## BACKGROUND

RNA structure is fundamental to understanding RNA function [1], as it undergoes dynamic changes in response to different cellular conditions. Accurate RNA structure analysis is therefore essential for uncovering these complexities. Techniques for determining RNA structure are broadly categorized into experimental and predictive methods. Experimental methods include traditional biophysical techniques such as X-ray crystallography, nuclear magnetic resonance spectroscopy, and cryo-electron microscopy [2], as well as chemical modification-based RNA structure probing methods utilizing high-throughput sequencing, such as DMS-seq, DMS-MaPseq, SHAPE-Seq, SHAPE-MaP, and icSHAPE [3–7]. Predictive methods encompass computational models, including machine learning and artificial intelligence frameworks like AlphaFold3 [8].

Chemical probing methods, compared to other approaches, provide the distinct advantage of detecting RNA structures *in vivo* and on a genome-wide scale [9]. Since 1980, chemical probing techniques have advanced significantly, with the DMS and SHAPE reagents being the most commonly used [1,10]. These reagents modify single-stranded RNA regions, which, during reverse transcription, result in either mutations in the complementary DNA (cDNA) or premature synthesis termination. The mutation and stop signals generated through this process provide indirect yet highly informative snapshots of RNA structure through high-throughput sequencing [11]. Current analysis methods, however, rely on only one primary signal (the signal used in the original method), neglecting the potential insights that could be gained from using both signals in tandem[12].

Novoa et al. used the support vector machines model to integrate primary stop signals from the DMS-seq experiment and primary mutation signals from the DMS-MaPseq experiment, improving accuracy by incorporating features like mutation direction ratios [13]. In addition, Sexton et al. found the existence of non-overlapping distributions of mutation and stop signals in mouse Xist lncRNA and 18S ribosomal RNA (rRNA) in one single experiment using DMS reagents, indicating that each signal type may independently carry structural information [11]. Therefore, integrating both signals from a single experiment may thus improve RNA structure signal analysis, but no studies have validated this approach on large-scale datasets [13,14].

Currently, most RNA structure datasets are derived from bulk RNA-Seq. With recent advancements in single-cell technology, it is now possible to analyze RNA structures at single-cell resolution. Wang et al. generated a single-cell RNA structurome dataset using a SHAPE-MaP-inspired method called sc-SPORT on the H9 and other cell lines [15]. In H9 cell line bulk datasets, accuracy for 18S rRNA typically reached approximately 0.69 across all nucleotides [16], whereas single-cell datasets achieved an accuracy of only 0.57-0.67 when low-accessibility nucleotides were filtered [15]. The observed discrepancy may stem from the inherently lower read coverage in single-cell data, a recognized limitation of single-cell RNA sequencing due to the minimal RNA input per cell [17]. Overcoming this limitation underscores the need for advanced methods to enhance RNA structure analysis accuracy in low-depth datasets, thereby enabling significant progress in single-cell RNA structurome research.

RNA-binding proteins (RBPs) and their interactions with RNA are crucial for understanding RNA function and regulation, playing key roles in processes such as splicing, translation, and RNA stability [18]. Current methods for detecting RBP binding sites primarily rely on techniques such as Cross-Linking Immunoprecipitation and sequencing (CLIP-seq) [19]. Studies have demonstrated that RNA structure is integral for RBP binding, and some predictive methods incorporating RNA structure have been developed [20]. Smola et al. non-targetedly explored the RBP binding sites of the mouse Xist lncRNA by comparing the differences in SHAPE scores between *in vitro* and *in vivo* conditions, and validated that the results were in high agreement with CLIP-seq data [21]. Although techniques such as LASER, which use NAz reagents to assess the exposure of RNA surface accessibility in solution, still depend on RNA structure probing experiments [22,23]. However, no methods rely solely on RNA structural signals from a single experiment to identify RBP binding sites [24].

In this study, we introduce a machine learning-based approach, STONE (Determine RNA **<u>S</u>**tructure with S**t**op-mutation Signals within **<u>ONE</u>** Experiment), which uses feature engineering to integrate mutation and stop signals from a single experiment, thereby enhancing signal analysis accuracy. We demonstrate the effectiveness of STONE in improving RNA structure analysis across genome-wide and single-cell datasets and in directly identifying RBP binding sites from a single RNA structure probing experiment.

## RESULTS

### Effectiveness and complementarity of mutation and stop signals in single experiment

Before developing the integration method, we systematically categorized the experimental workflows into two groups: 1) those capable of generating both signal types (**Figure S1A**), and 2) those that generate only one type of signal (**Figure S1B**). For datasets containing both signal types, we aimed to validate that both signals co-exist and contain effective structural information. We designate the signal used in the original method as the primary signal (e.g., stop signal in DMS-seq), while the supplementary signal is referred to as the secondary signal (e.g., mutation signal in DMS-seq) (**Figure S1C** shows all methods and their primary and secondary signal).

Using double-stranded regions of known RNA structures as controls, we confirmed that the signal in these regions is significantly lower than in single-stranded regions, indicating the validity of both primary and secondary signals (**Figures 1A-B**). We also validated the efficacy of secondary signals for RNA structure analysis using a larger dataset with various modification methods, including unmodified DMSO groups as controls. Our analysis showed that secondary signals from the modified group had significantly higher accuracy compared to both primary and secondary signals from the DMSO control (**Figure 1C**), confirming secondary signals provide valuable structural information for RNA structural analysis. Another key piece of evidence is base bias. The DMS reagent preferentially modifies adenine (A) and cytosine (C) bases, which we observed in both the primary and secondary signals of DMS-seq and DMS-MaPseq (**Figures 1D-G**), confirming that the secondary signals are generated from DMS reagent modification events. Furthermore, for SHAPE methods, the mutation signal from the secondary signals of icSHAPE strongly correlated with the mutation signal from the primary signals of sc-SPORT (Pearson correlation r = 0.756 ± 0.040, **Figures S1D-E**), confirming their structural relevance.

**Figure 1.**
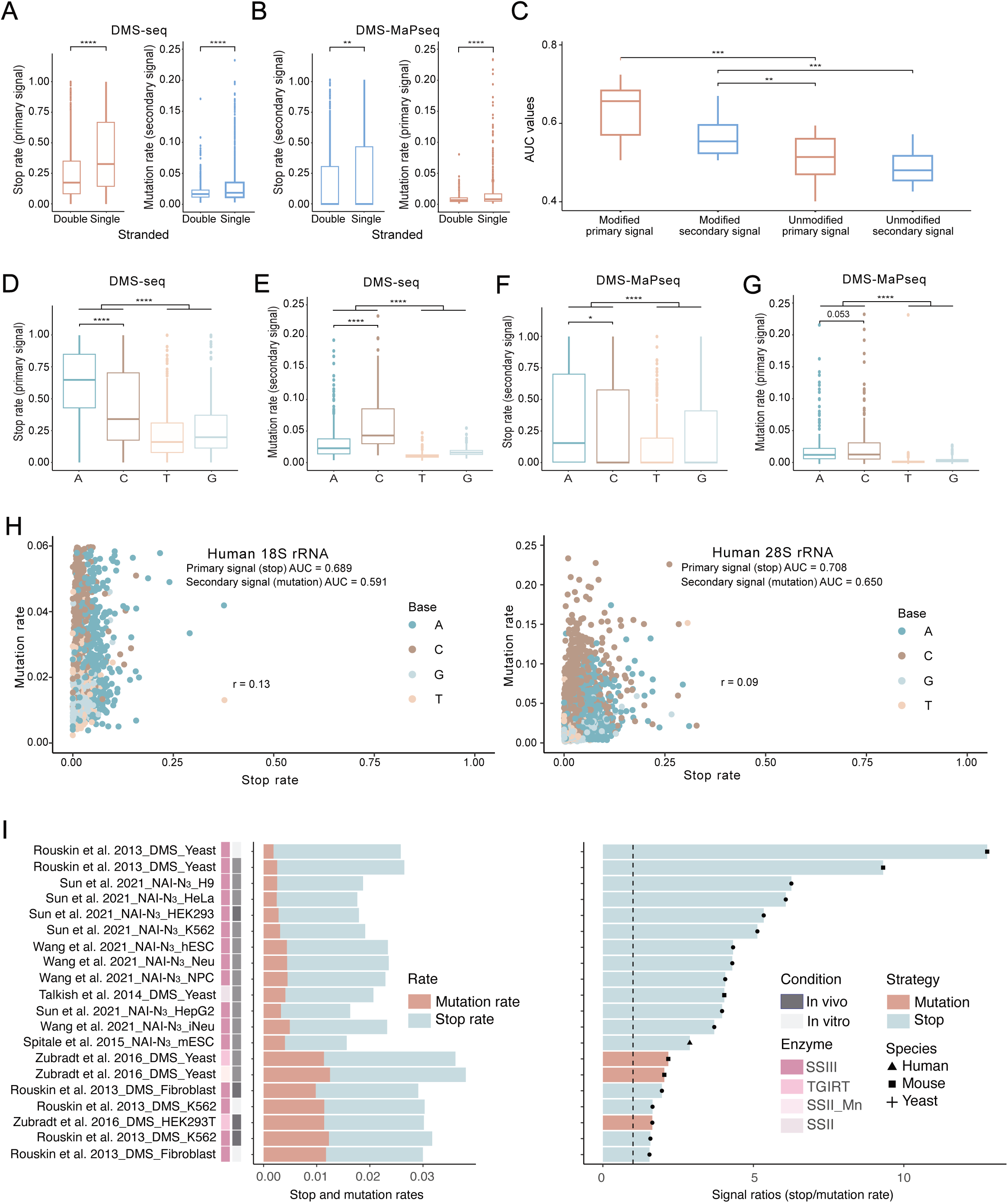
Summary of signal compositions for RNA structure probing protocols. (A-B) Comparison of primary and secondary signal distributions in single-stranded and double-stranded regions of human 18S rRNA. The levels of significance displayed in the plots were calculated using unpaired t-tests. (C) Accuracies of the original primary and secondary signals for both small molecules modified and DMSO control groups. Datasets (n=23) included results from DMS-seq, Structure-seq, DMS-MaPseq, SHAPE-MaP, and icSHAPE protocols. The levels of significance displayed in the plots were calculated using paired t-tests. (D-G) Base biase preference analysis of chemical modification signals. All points are from the single-stranded regions of human 18S rRNA. The levels of significance displayed in the plots were calculated using unpaired t-tests. (H) Scatter plots showing the correlation between primary stop and secondary mutation signals in human 18S and 28S rRNA in DMS-seq dataset [26]. (I) Bar plots summarized total signal rates (left) and signal ratios (right, calculated as stop rate/mutation rate) from genome-wide RNA structure probing datasets. Data were compiled across multiple publications involving different enzyme and reagent combinations. The dashed vertical line (right) was plotted at signal ratio=1. Statistical significance is denoted as follows: n.s. (not significant) for p > 0.05, * for p < 0.05, ** for p < 0.01, *** for p < 0.001, **** for p < 0.0001.

According to previous research, although both types of signals can be detected in a single experiment, their distributions show low correlation[11]. Similarly, we observed low base distribution correlations between the base distributions of the mutation and stop signals in human, yeast, and virus datasets (**Figures 1H** and **S1F-G**). This indicates that they capture complementary structural insights. Furthermore, despite the low signal distribution correlations observed between primary and secondary signals (**Figure 1H**), we also found complementary signal distribution patterns (**Figures 1D-G**): specifically, stop signals—whether from the primary signal in DMS-seq or the secondary signal in DMS-MaPseq—tend to favor A bases (**Figures 1D** and **1F**), while mutation signals show a preference for C bases (**Figures 1E** and **1G**). This observation suggests that relying on only one signal type in an experiment could lead to the loss of distributional characteristics inherent in the other signal type.

Finally, we performed a comprehensive dual-signal quantification across 20 datasets from 6 studies (**Table S1**). At the genome-wide level, the two-signal phenomenon was also observed. Notably, even in mutation-centric strategies, most methodologies generated two types of signals, largely consisting of stop signals (**Figure 1I**). This result aligns with the understanding that not all stop signals are due to chemical modifications, some arise simply from sequence fragmentation [25]. To investigate signal distribution during technological development, we focused on yeast as a model organism. DMS-seq, primarily driven by stop signals, exhibited a low mutation rate of 0.19-0.26% (**Figure 1I**). Three years later, DMS-MaPseq was introduced, which increased mutation rates in yeast to 1.14-1.25% by utilizing new reverse transcriptase TGIRT (**Figure 1I**). However, even before the advent of DMS-MaPseq, higher mutation rates were already observed in DMS-seq datasets for mammalian cells. For example, the mutation rate of DMS-seq reached 1.23% in the Fibroblast and K562 cell line datasets [26]. This is almost consistent with the 1.14% mutation rate of DMS-MaPseq in the HEK293T cell line samples [27]. This suggests that the potential for integrating two signals was already present in the early stages of DMS-seq technology development.

### Signal features extraction and integration method development

Based on the evidence supporting the effectiveness and complementarity of the two signal types, we developed a method called STONE, that integrated two signals using feature engineering and machine learning models (**Figure 2**). As shown in **Figure 2**, the workflow involved separately analyzing the mutation and stop signals. Then, feature engineering was used to expand the features of these signals, which were then applied to a machine learning model to calculate the probability of each nucleotide being single-stranded, known as the STONE scores.

**Figure 2.**
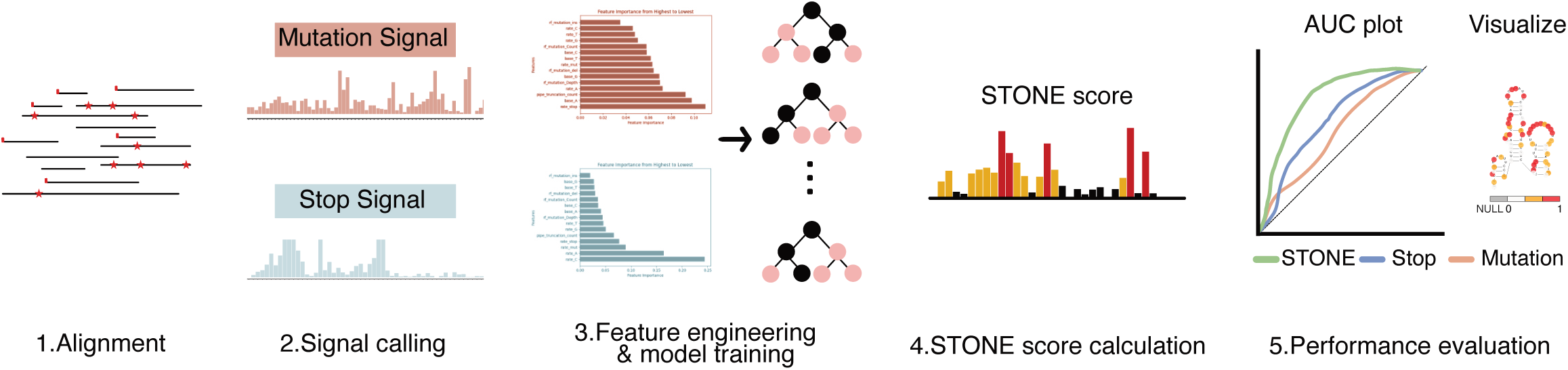
STONE method workflow. The workflow illustrates the key steps involved in RNA structure signal calling and model training for STONE method.

Traditional methods rely solely on counts and base depths of primary signals, failing to fully utilize the rich information potentially contained in base types. A comprehensive evaluation of STONE was conducted across five machine learning models (**Figure S2A**) and nine deep learning models (**Figure S2B**), utilizing up to 42 features (**Table S2**), such as the relative proportion of each base in the total sequencing depth. To balance learning capacity and generalization while maintaining simplicity and minimal parameter usage, the optimal performance in machine learning methods was achieved by the Random Forest model, which outperformed all deep learning models in this simple parameter task (**Figures S2A-B**). Fifteen key features were ultimately selected for training the signal integration model using a Random Forest model (**Figure S2C**). Due to substantial differences in chemical modification mechanisms, we divided the training into two models: one for SHAPE reagents and one for DMS reagents. The models were trained only using 18S rRNA. For parameter tuning, 25% of the nucleotide sites, including both positive (unpaired) and negative (paired) sites, from each dataset were reserved as the validation set.

As shown in **Figure S2C**, in the DMS model, the main contributing features were C and A base rates, while the SHAPE model has numerous features contributing uniformly. This precisely aligns with the understanding that DMS primarily targets C and A bases, while SHAPE can modify all four types of bases unbiasedly [1]. Specifically, the contributions of various types of parameters to the final AUC values differ among different strategies, but all features contributed effectively to elevate accuracies than using only traditional observed features (**Figure S2D**). As shown in **Figure S2E**, the integrated signals of STONE effectively combine complementary information from dual signals.

### Performance validation of STONE across probing strategies

The accuracy of the STONE method was evaluated by comparing STONE scores to known RNA structures on a single-gene scale. Its performance was assessed across the entire transcript, within specific structural motifs, and at individual nucleotide positions.

Notably, the two models were trained exclusively on 18S rRNA datasets from various studies (**Table S3**). The validation set consisted of 25% of the initial 18S rRNA nucleotides, ensuring an unbiased internal evaluation of the model during training. The test set included additional 18S rRNA datasets from external studies, as well as other RNAs with well-characterized structures, demonstrating that the STONE method effectively improved accuracy (**Figures 3A** and **S3A**). Additionally, we observed that when the AUC derived from original signals is relatively low, as observed in RNAs such as human SRP RNA and U1 snRNA, the improvement in accuracy provided by STONE remains modest (**Figure 3A**). Conversely, for RNAs like human 28S rRNA, where the AUC for both signals is higher, the integration of signals by STONE yielded a significant improvement, with the final AUC reaching as high as 0.849 (**Figure 3A**). This demonstrates that STONE consistently enhances AUC values relative to the original scores, though datasets with effective dual signals are necessary to ensure optimal model performance.

**Figure 3.**
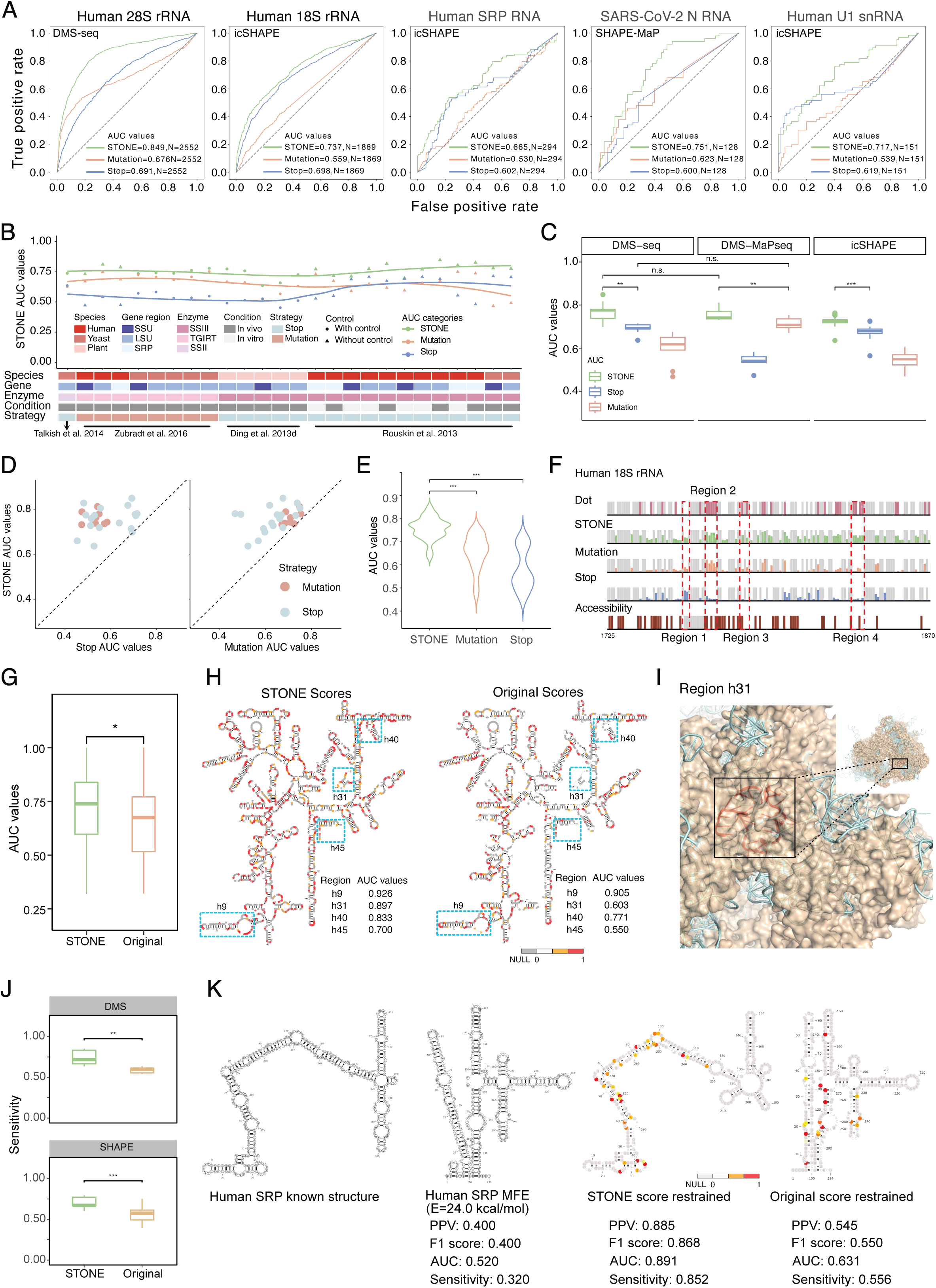
Model performance evaluation and validation across RNA structure probing datasets. (A) Comparison of AUC values for the original and STONE applied to human 18S rRNA, human 28S rRNA, human SRP RNA, human U1 snRNA, and SARS-CoV-2 Nucleocapsid protein (N) mRNA. (B) Validation of DMS model performance via AUC values across datasets from different species, gene regions, experimental conditions, small molecule-enzyme combinations, and reverse transcription strategies. Paired t-tests were used to calculate p-values. (C) Evaluation of model performance across different RNA structure probing workflows using box plot. Paired t-tests were employed to compare AUC values within the same experimental method, while unpaired t-tests were applied to compare AUC values across different methods. The numbers of datasets used are as follows: n=9 for DMS-seq, n=7 for DMS-MaPseq, and n=40 for icSHAPE. (D, E) Statistical analysis of AUC values from 26 datasets, providing a comprehensive evaluation of the DMS model across various experimental contexts. The p-values were calculated using paired t-tests. (F) Single-nucleotide resolution improvements in RNA structural scores for human 18S rRNA, published by Zubradt et al. [4] Results were visualized by IGV (Integrative Genomics Viewer). The red boxes highlighted some examples of successive positions with evident improvements. (G) The accuracies of structural scores for 45 regions of human 18S rRNA were compared between STONE and the traditional data analysis pipeline. Statistical significance was calculated using unpaired t-tests, and the corresponding p-values are displayed in the plots. (H) Visualization of the accuracy and coverage of structural scores derived from the original and STONE on the established 18S rRNA secondary structure (obtained from the RNA STRAND database [50]). AUC values for regions highlighted by blue boxes were independently calculated and annotated on the plot. Nucleotides were colored using structural scores. (I) Visualization of RNA and RBP structure and interactions in region 2 of the human 18S rRNA from plot G (colored in red). The structural model of human 18S rRNA was obtained from the Protein Data Bank (PDB ID: 4V6X). RNA was colored in cyan, and protein was colored in wheat. (J) Evaluation of the accuracy of structural predictions for DMS and SHAPE datasets under sensitivity metrics. The box plots were derived from 30 human datasets, containing various experimental methods (DMS-seq, DMS-MaPseq, icSHAPE, SHAPE-MaP) and distinct gene regions (18S, 28S, SRP, U1). Predicted structures obtained using STONE analysis and those derived from traditional methods were compared to known structures. Statistical significance was assessed using paired t-tests, with corresponding p-values displayed in the plots. (K) The folded SRP RNA structure under various constraints. The resulting structures were evaluated using several performance metrics: PPV, AUC, F1-score, and sensitivity. These metrics are used to assess the accuracy of RNA structure predictions derived from both the STONE method and the original folding method, with respect to a known reference structure. Statistical significance is denoted as follows: n.s. (not significant) for p > 0.05, * for p < 0.05, ** for p < 0.01, *** for p < 0.001, **** for p < 0.0001.

Independent test datasets were employed to evaluate model performance, as outlined in the methods section. Improvements in accuracy were widely observed across well-characterized structures in both DMS and SHAPE datasets (**Figures 3B** and **S3B**). As previously mentioned, when DMS-seq first appeared in 2013, there was a high proportion of both signals present in human cell line samples. Intriguingly, STONE’s integration of these signals in DMS-seq displayed a trend toward higher accuracy than that observed with the DMS-MaPseq method introduced in 2016, underscoring the potential benefits of signal integration for enhanced RNA structure analysis (**Figure 3B**).

Overall, whether employing DMS-type or SHAPE-type strategies, both stop and mutation approaches led to significant improvements in accuracy (**Figures 3D-E** and **S3C**). To confirm that the widespread improvement was not due to overfitting, we masked the secondary signals, leading to a significant performance reduction (**Figure S4**). This confirms that STONE relies on the combined contributions of both primary and secondary signals. Furthermore, the random shuffling of secondary signals further validated that their positional correspondence is crucial, as randomized integration failed to improve accuracy (**Figure S5**). This highlights that STONE requires authentic paired two types of signal features for successful integration and does not function effectively with arbitrary input.

We evaluated the STONE method across various modification strategies (**Figure 3C**) and found that it consistently improves accuracy, performing well across different modification intensities and reverse transcriptase conditions (**Figures S6A-C**). STONE enhanced performance for both mutation-based and stop-based reverse transcriptases, and showed broad applicability with various chemical modification reagents, even without retraining for some reagents like 1M7 and NAI (**Figure S6D**). This suggests the model generalizes well across RNA structure probing experiments, likely due to the similar structural effects of small molecule modifications on the reverse transcription process (**Figure S6F**), aligning with their respective modification mechanisms [28].

The nucleotide resolution results, exemplified by human cell lines using DMS-MaPseq (**Figure 3F**) and icSHAPE (**Figure S3D**) methodology, showed improved accuracy in four representative regions. As demonstrated in **Figure 3F**, the STONE method could decrease the signal intensity in double-stranded regions with existing signals (Region 1), increase the signal intensity in single-stranded regions with low signals (Regions 3 and 4), and resolve ambiguity in regions with conflicting signals by accurately identifying more reliable information at nucleotide resolution (Region 2).

In the region resolution validation, human 18S rRNA was subdivided into 45 defined regions [29] (**Figure S7A**), and we observed an universal AUC improvement with STONE compared to the original method across these regions (**Figures 3G** and **S7B**). Notably, the four regions highlighted in **Figures 3H** exhibit varied AUC increases, including 0.021, 0.294, 0.062, and 0.150, corresponding to improvements from 0.905 to 0.926, 0.603 to 0.897, 0.771 to 0.833, and 0.550 to 0.700, respectively. In particular, the region h31 shows the most pronounced enhancement, likely due to its central location amid ribosomal proteins that limit small molecule accessibility, resulting in low signals that are challenging to resolve with traditional methods (**Figure 3I**). The region h31 also showed improvement using the SHAPE model (**Figure S7C**).

The region h45 (1838-1870) is located in the central zone, where intricate biomolecular interactions, such as ribosome and mRNA binding, occur [15]. These complex interactions make it challenging to obtain accurate chemical modification information. Surprisingly, the STONE model still demonstrated high fidelity in improving RNA structure probing accuracy within this complicated region, highlighting its broader potential for application (**Figure S8**).

To assess the utility of dual-signal integration in improving RNA structure prediction, we first optimized the folding parameters for STONE (Slope = 3.0 kcal/mol, Intercept = −0.6 kcal/mol) and traditional methods (Slope = 3.4 kcal/mol, Intercept = −1.4 kcal/mol) on 18S rRNA (**Figure S9A**) and compared their sensitivity to confirm alignment with known structures. We found that the STONE method (Sensitivity = 0.775 ± 0.048) showed significantly higher alignment with the known structure compared to the original method (Sensitivity = 0.522 ± 0.079) (paired t-test p = 2.3 × 10^-5^, **Figure S9B**). We further expanded the dataset and observed gene regions to 10 datasets with 2 gene regions (paired t-test p_18S_ = 2.3 × 10^-6^, p_28S_ = 5.5 × 10^-6^, **Figure S9C**) and 22 datasets with 4 gene regions (paired t-test p_DMS model_ = 2.0 × 10^-3^, p_SHAPE model_ = 1.8 × 10^-4^, **Figures 3J** and **S9D**), and consistently found that RNA structures predicted using the STONE score exhibit significantly higher similarity to the reference structures. Among the 18S, 28S, SRP, and U1 gene regions, we used SRP as an example to visually illustrate the contribution of the STONE score. The STONE-constrained structure for SRP RNA closely matched the known reference, highlighting the improved performance of STONE scores in RNA folding prediction (**Figures S9E** and **3K**). These results indicate that integrating mutation and stop signals via the STONE method not only enhances RNA structural signal quality but also improves RNA folding predictions, demonstrating its value in RNA structure modeling.

### STONE enhances accuracy and coverage in genome-wide analysis

To assess the broader applicability of the STONE method for RNA structure studies, we conducted genome-wide analyses. During the data collection, we evaluated the sequencing depth of genome-wide datasets, observing that as the sequencing depth requirements increased, the number of detectable genes correspondingly declined (**Figure 4A**). This observation emphasized the need to evaluate the STONE’s performance under conditions of reduced sequencing depth. Through downsampling experiments on 18S and 28S rRNA, we found that the STONE method maintained significantly higher accuracy than traditional approaches even at low sequencing depth (paired t-test, p < 0.001, **Figures 4B** and **S10**), underscoring its potential for genome-wide application.

**Figure 4.**
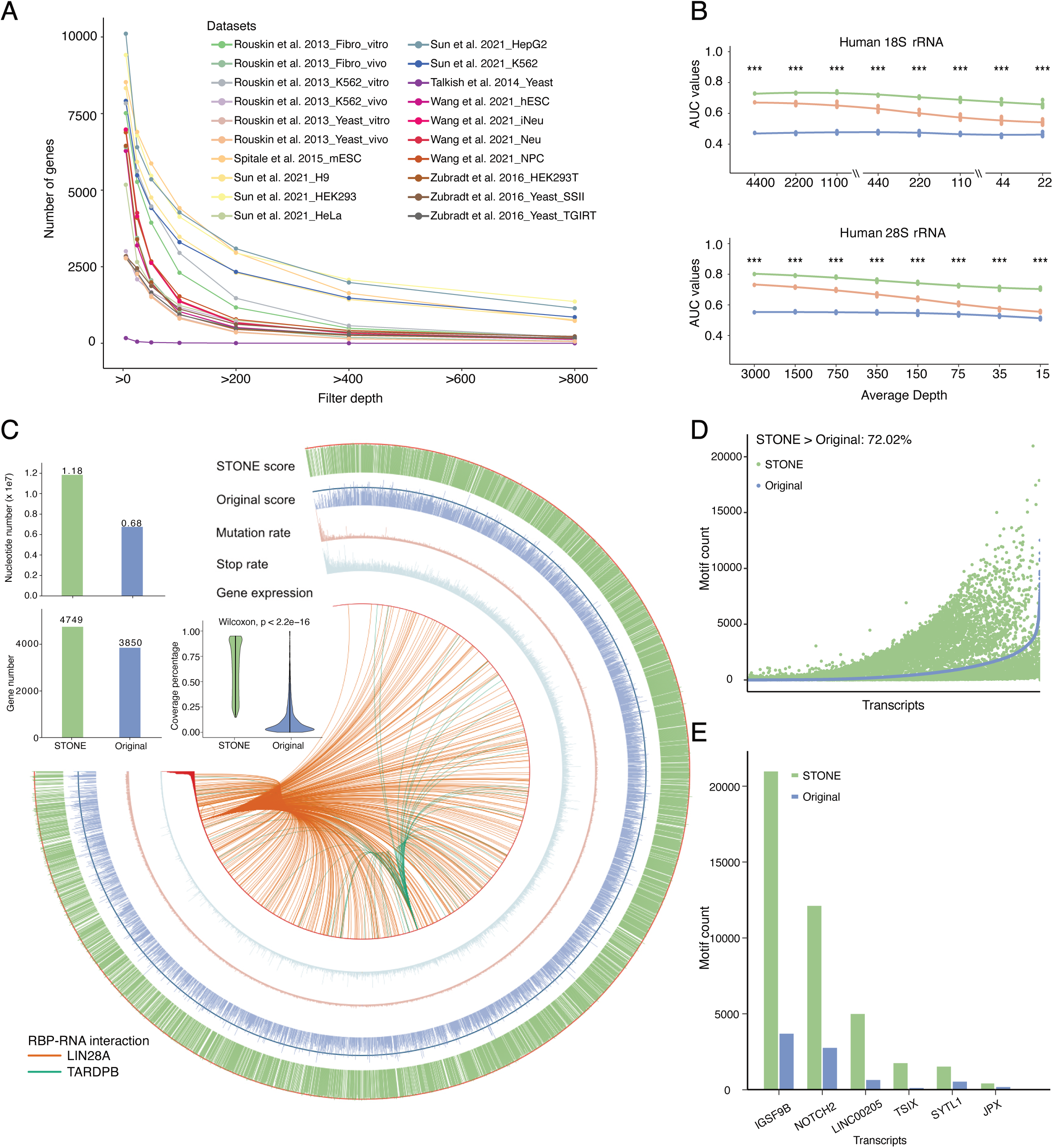
Evaluating STONE performance when applying to genome-wide RNA datasets. (A) line plot illustrating the number of genes covered with effective signals after filtering genome-wide datasets based on various depth thresholds. (B) Evaluation of STONE performance using down-sampled datasets reported by Zubradt et al. [27], conducted with DMS-MaPseq. The p-values were calculated using paired t-tests to assess the significance of differences between STONE and original calculations (n=5). (C) Global analysis of the genome-wide H9 dataset from Sun et al. [16], conducted using icSHAPE. The arc plot was used to display signal distribution and structural scores for all H9 transcripts. Effective signals were summarized by gene number, nucleotide number, and transcript coverage percentage. The p-value was calculated using the Wilcoxon test to compare the significance of differences in coverage percentages. Curved lines within the circular plot represent RNA-RBP interactions, with orange lines depicting LIN28A-RNA interactions, and green lines depicting TARDPB-RNA interactions. (D) Dot plot comparing the number of stem-loop structures identified using STONE-derived versus original structural scores across all transcripts with valid signals. The percentage represents the proportion of identified transcripts possessing more stem loops when analyzed using STONE compared with the original stop signal-based calculations. (E) Different types of RNAs with high, median, and low motif counts derived from STONE and original scores. Statistical significance is denoted as follows: n.s. (not significant) for p > 0.05, * for p < 0.05, ** for p < 0.01, *** for p < 0.001, **** for p < 0.0001.

To evaluate the improvement in gene detection and nucleotide coverage with STONE (details in Methods, **Figure S11**), we applied an effective sequencing depth threshold—defined as the minimum depth at which the STONE AUC declines to 80% of its maximum value observed in the undownsampled dataset—to our genome-wide analysis of the H9 cell line icSHAPE dataset [16]. We observed a 23.4% increase in gene detection and a 75.3% increase in nucleotide coverage, thereby substantially enhancing the availability of robust structural signals across transcripts (paired Wilcoxon test, p < 0.001, **Figure 4C**). The genome-wide distribution of both mutation and stop signals was visualized using the longest transcripts as references (**Figure 4C**). Structural scores derived from the original calculation indicated that highly expressed transcripts exhibited higher structural scores, with a greater proportion exceeding the 75% threshold (**Figure 4C**). This pattern was also observed in the STONE-generated structure scores, aligning with previous studies suggesting a positive correlation between RNA structural unpairing and transcriptional activity [30].

Given the functional significance of structural motifs like the stem loops, we further examined the STONE method’s capability to identify more functionally relevant motifs compared to traditional approaches. The structural motif searches targeted stem loops with stem lengths ranging from 2 to 7 nt and loop lengths between 2 and 7 nt [31]. In the H9 cell line dataset, the STONE method identified more structural motifs in 72.0% of all transcripts (**Figure 4D**). We selected six representative transcripts based on their structural motif counts (**Figure 4E**): IGSF9B and NOTCH2 RNA with high counts, LINC00205 and TSIX RNA with medium counts, and SYTL1 and JPX RNA with low counts. For the TSIX, JPX, and LINC00205 lncRNAs, the STONE method detected increases of 15.4 folds, 2.5 folds, and 7.7 folds, respectively, in structural motif numbers. Similarly, the protein-coding IGSF9B, NOTCH2, and SYTL1 mRNA showed increases of 5.7 folds, 4.4 folds, and 2.8 folds, respectively. These results underscore STONE’s effectiveness in revealing critical structural motifs from refined structural scores, which may contribute to the discovery of underlying gene regulatory mechanisms.

### STONE enhances single-cell analysis with superior accuracy and efficiency

The advancement of single-cell technologies has revolutionized RNA structure analysis, enabling probing at single-cell resolution [15]. However, current single-cell technologies still rely on DMSO treatment as background control to improve the signal-to-noise ratio. By validating STONE on bulk datasets, we demonstrated that STONE’s superior accuracy is not limited by the absence of control information (**Figure 5A**). As displayed in the boxplot, the STONE method demonstrated superior accuracy on bulk datasets, achieving an average AUC of 0.74 compared to 0.69 for traditional control-dependent methods, while still maintaining robust performance at 0.72 even without control data (**Figure 5A**). This finding suggests that the STONE method holds significant promise for single-cell RNA structure analysis, as it overcomes the limitations of single-cell experiments where it is nearly impossible to establish a perfect control group.

**Figure 5.**
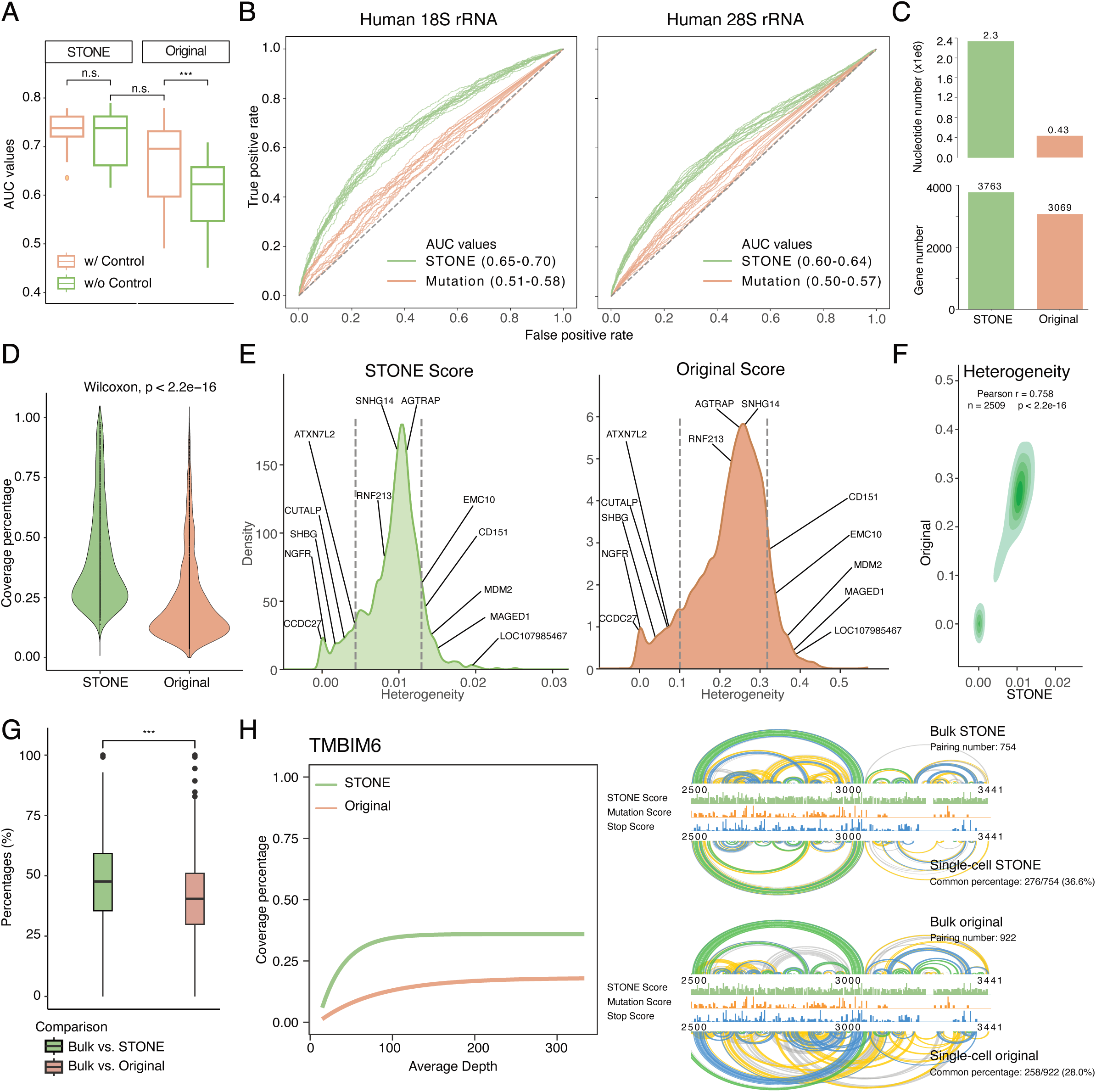
Performances evaluation of STONE in single-cell RNA structure analysis. (A) Box plots comparing model performance with and without the inclusion control groups. Paired t-tests were employed to compare AUC values obtained from the same calculation method, while unpaired t-tests were applied when comparing AUC values across different calculations (n=12). (B) Plot of AUC values derived from STONE versus original AUC values, using data from 15 cells reported by Wang et al. [15]. (C) Comparison of effective signal capture at both gene and nucleotide levels using STONE calculation versus the original calculation. (D) The signal coverage percentage for each identified transcript was compared between STONE and original calculation methods. The p-value was calculated using the Wilcoxon test, assessing the significance of the increased coverages. (E) Heterogeneity analysis for 76 cells from the H9 cell line using STONE and original-derived structural scores displayed in density plots. (F) Pearson correlation coefficient calculated to compare gene heterogeneity values derived from STONE and original structure scores across all available transcripts. (G) Structural similarity comparison of RNA structures (n=653) derived from bulk and single-cell datasets using different calculation methods. Paired t-tests were employed to examine improvements in similarity percentages when using STONE. (H) TMBIM6 transcript was used as an example to show similarity between RNA structures derived from bulk and single-cell datasets. The colors of the arc represent the confidence level of the corresponding pairing, ranked from highest to lowest as follows: green (80%-100%), blue (30%-80%), yellow (10%-30%), and grey (3%-10%). Statistical significance is denoted as follows: n.s. (not significant) for p > 0.05, * for p < 0.05, ** for p < 0.01, *** for p < 0.001, **** for p < 0.0001.

To validate the performance of STONE, we adopted Wang et al.’s single-cell dataset using the sc-SPORT approach [15]. In the original study, accessibility information for 18S rRNA was used to exclude regions with low accessibility from the AUC calculation, yielding AUC values of 0.57-0.67 for 18S rRNA and 0.49-0.58 for 28S rRNA when both control and accessibility data were included (**Figure S12 left**). Under these same conditions, STONE achieved higher AUC ranges: 0.66-0.72 for 18S and 0.58-0.62 for 28S rRNA (**Figure S12 left**). Notably, when accessibility information was excluded, STONE’s AUC for well-characterized 18S (0.65-0.70) and 28S rRNA (0.60-0.64) regions was only minimally affected, whereas the original method’s AUC values dropped significantly to 0.51-0.58 and 0.50-0.57, respectively (**Figure 5B**). With both control and accessibility data removed, the original method’s accuracy fell to about 0.5, indicating unreliable performance (**Figure S12 right**). In contrast, STONE maintained consistently high AUC values, with 18S rRNA ranging from 0.62 to 0.65 and 28S rRNA from 0.59 to 0.62 (**Figure S12 right**). Collectively, these results demonstrate STONE’s robustness in preserving high accuracy without reliance on control or accessibility information, underscoring its suitability for single-cell RNA structure analysis.

To further investigate STONE’s suitability for single-cell RNA structure analysis, we assessed its ability to expand RNA structure signal coverage under stringent depth thresholds, defined by the minimum AUC-supporting depth for reliable detection. The STONE method uncovered 27.6% more genes and 431.7% more nucleotides (**Figure 5C**). Focusing on individual transcripts, STONE significantly enhanced signal coverage (paired Wilcoxon test, p < 0.001, **Figure 5D**). This enhanced signal coverage also reduced undesired variations among individual cells, resulting in more consistent transcript structures compared to traditional approaches (**Figure 5E**). To be specific, heterogeneity scores for individual genes were reduced, ranging from 0 to 0.036 compared to 0 to 0.568 using traditional methods (**Figure 5E**). Importantly, despite this reduction in heterogeneity, the relative positioning of transcripts remained stable, indicating that meaningful heterogeneity information could still be accurately extracted from STONE-derived structural scores (Pearson correlation r = 0.76, **Figure 5F**).

STONE improved the reliability of single-cell RNA structures by increasing transcriptomic base-level coverage. Due to STONE’s high efficiency in extracting signals, especially in low-depth data, we assessed the signal reliability by comparing the single-cell RNA structures generated by STONE to those derived from high-depth datasets. Overall, STONE significantly improved the number of common pairs with bulk data (paired t-test, p < 0.001, **Figure 5G**). Compared to bulk datasets, STONE in the single-cell dataset improved structural consistency in 67.9% of transcripts, with average improvement rates (26.1%) exceeding average reduction rates (7.3%) by more than threefold. For instance, STONE provided broader signal coverage for LIN28A-targeted RNA TMBIM6 compared to the original method, achieving saturation coverage with fewer reads (**Figure 5H**). We compared the similarity of TMBIM6 structures by quantifying the number of common base pairs between single-cell and bulk structures for both STONE and the original method. **Figure 5H** shows that STONE produced more consistent pairings (36.6%) for single-cell TMBIM6 structures compared to the original calculations (28%) in the covered region of TMBIM6 (positions 2500-3441). Specifically, around position 3100 of TMBIM6 RNA, the original method, relying only on mutation signals, showed numerous low-confidence, long-distance pairings (**Figure 5H**). By integrating both mutation and stop signals, STONE enhanced signal intensity in this region, reducing false pairings and improving the overall reliability of the single-cell TMBIM6 structure (**Figure 5H**).

### STONE directly identifies RBP binding sites from RNA structure probing

In analyses of well-characterized structures such as 18S rRNA, specific regions show markedly reduced signal intensities, primarily due to the extensive RNA-protein interactions that restrict the accessibility of chemical modification agents [26]. While masking low-accessibility regions can enhance RNA structure analysis accuracy (**Figure S12**), comprehensive accessibility data remain unavailable for most RNAs. Therefore, we investigated the dependence of the STONE method on accessibility information. In 18S rRNA studies where small-molecule accessibility information was excluded, the decrease in AUC for the STONE method was smaller than that for the traditional method (ΔAUC_STONE_ = 0.049, ΔAUC_original_ = 0.082, **Figure 6A**). This indicates that the STONE method performs better at preserving accuracy in regions with lower accessibility. Moreover, compared to the original method, STONE consistently exhibited significantly higher AUC without accessibility (paired t-test, p = 7.92 × 10^-11^, **Figure 6A**). This indicates that the STONE method is less dependent on accessibility information.

**Figure 6.**
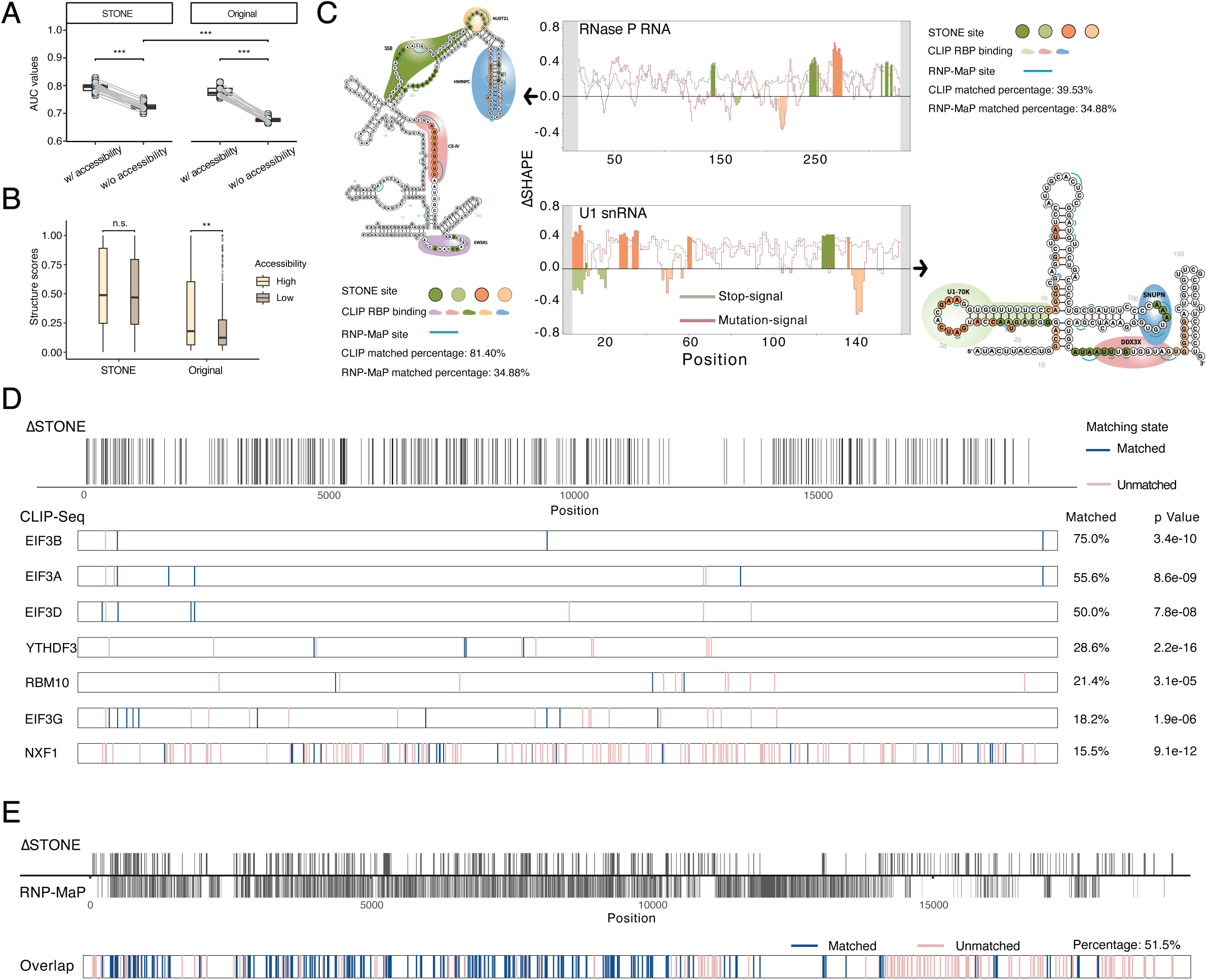
Utilizing STONE-derived structural scores to identify RNA-RBP interactions. (A) Model performance comparison on 18S rRNA with and without protein accessibility information. Paired t-tests were employed to compare AUC values between methods. The number of datasets used is as follow: n=38 for NAI-N_3_. (B) STONE performance on identifying the single-strand positions on human 18S rRNA without providing the accessibility information. Paired t-tests were applied to calculate p-values. The number of positions used are as follows: n=175 for high accessibility and n=769 for low accessibility. (C) ΔSTONE scores were calculated using STONE-derived structural scores to identify RBP binding locations on human U1 snRNA and human RNaseP (based on the H9 dataset from Sun et al. [16]). The calculated positions were visualized using known RNA structures from the RNA Central database. ΔSTONE outcomes (colored nucleotides) were compared with CLIP-seq (demonstrated by drawing RBPs) and RNP-MaP (displayed using blue lines) results and the matching percentages for each of the comparisons were annotated. (D) ΔSTONE-calculated RBP binding data on human XIST lncRNA were aligned with CLIP-seq data at the regional level, the level of alignment was compared to randomly sampled regions for each RBP. t-tests were performed using 10 random matching percentages and the ΔSTONE versus CLIP-seq matching percentage for each RBP. (E) Comparison of RBP binding regions on human XIST lncRNA calculated using ΔSTONE versus RNP-MaP. Statistical significance is denoted as follows: n.s. (not significant) for p > 0.05, * for p < 0.05, ** for p < 0.01, *** for p < 0.001, **** for p < 0.0001.

Next, we analyzed the main contributing factor to the improvement in accuracy of the STONE method in low-accessibility regions (spatial distance between protein and RNA < 3 Å) [32]. In single-stranded regions with low accessibility, the structural scores of the STONE method were not significantly impacted compared to those in high-accessibility regions (ΔScore_STONE_ = 0.028, p = 0.317, **Figure 6B**). In contrast, the original method showed a significant decrease in structural scores under the same conditions (ΔScore_original_ = 0.093, p = 0.002, **Figure 6B**). This observation highlights that comparing regions with significant differential signal profiles to traditional methods (ΔSTONE) derived from a single RNA structure probing experiment could facilitate the identification of RNA regions impacted by protein binding.

To further validate the efficacy of the STONE method, we evaluated its performance on both short RNAs, such as U1 snRNA and RNase P RNA, as well as long RNAs like XIST lncRNA. These results were then compared with those obtained from antibody-dependent CLIP-seq techniques and the non-specific RNP-MaP approach. RNP-MaP is a live-cell chemical probing strategy that uses a hetero-bifunctional crosslinker to map RNA-protein interaction networks at nucleotide resolution by identifying multiple proteins bound to single RNA molecules through reverse transcription and sequencing [33]. For U1 snRNA and RNase P RNA, significate differential nucleotides identified by ΔSTONE showed strong alignment with CLIP-seq results (39.5% and 81.4% respectively) and RNP-MaP results (34.9%) (**Figure 6C**). For the long RNA XIST lncRNA, an evaluation using 33 CLIP-seq datasets in the HEK293 cell line revealed that STONE identified interactions for the majority of RBPs analyzed. It achieved overlap rates exceeding 15% for 31 RBPs, with 7 of these showing statistical significance (**Figure 6D**). Also, despite CLIP-seq experiments, the STONE method achieved a consistency rate of 51.5% when compared to non-targeted RBP detection techniques like RNP-MaP (**Figure 6E**). This consistency aligns with overlap rates commonly observed across experimental techniques, between different cell lines, or between experimental results and AI-based predictions, which typically range from 21.5% to 56.0% [20].

The STONE method’s ability to enhance signal intensity at nucleotides affected by protein interactions highlights its potential as a versatile tool for investigating RNA-protein interactions, particularly in regions where traditional methods face limitations.

## DISCUSSION

In traditional RNA chemical modification signal analysis, methods have typically relied on primary signals and a limited set of features. These methods involve acquiring signal counts and site depth features from alignment results, and using simple formulas for structural score calculation[25,34]. Previous studies have demonstrated the potential for integrating dual signals by analyzing base distribution correlations [11] and combining data from different modification strategies [13]. In addition, we demonstrated secondary mutation signals are present in stop-based data (**Figures S1D-E**). The signal rates in single- and double-stranded regions (**Figures 1A-B**) and base biases preferences in mutation and stop reaction (**Figure 1D-G**) further validate the authenticity of these signals. Therefore, based on their efficacy (**Figures 1A-C** and **S1D-E**) and complementarity (**Figures 1D-H**), we conclude that integrating both signals in RNA structure analysis is methodologically feasible. We introduce the STONE method, which enhances RNA structural signal analysis by integrating mutation and stop signals obtained from a single experiment using 15 expanded features (**Figure 2**). Using extensive datasets (**Table S1**), we validated the DMS and SHAPE models of STONE at well-characterized RNAs, demonstrating improved accuracy over traditional methods (**Figure 3**).

Our comprehensive evaluation revealed that datasets using stop strategies offered the most substantial improvements in accuracy with STONE (**Figure S6C**), as these strategies typically include high distributions of both mutation and stop signals (**Figure 1I**). Since the expanded base-specific features were mainly derived from mutation signals, leveraging mutation-related features provided additional support to the stop signals, enhancing overall AUC values (**Figure S2C**). We also found that single-cell RNA structurome analysis is particularly suited to STONE, as low coverage often limits traditional methods, and single-cell data lack reliable control groups. Currently, the availability of single-cell probing datasets is limited, with only one based on the SHAPE-MaP strategy [35]. The performance of STONE with other stop-centric strategies remains unexplored. According to the current evaluation, STONE’s performance with stop-centric strategies is expected to yield superior results (**Figure S6C**). Overall, we observed that STONE provided the more improvements in datasets with lower accuracy for the original primary signals. (**Figure S6E**).

STONE’s ability to learn accessibility information makes it valuable for applications like RBP binding site identification (**Figure 6A**). RNA structure analysis *in vivo* is challenging due to the complex, dynamic RNA folding and extensive RNA-protein interactions [36]. The likelihood of effective chemical modification varies in regions tightly bound by RBPs due to differences in the half-lives of modification reagents [28]. Smola et al. suggested that 1M7 works well for *in vivo* and *in vitro* comparisons to identify RBP binding sites, while NAI, due to its longer half-life, may be less suitable for capturing RBP-related signals [21]. Corley et al. also found DMS is more effective than NAI for identifying RBP binding information [37]. However, recent studies indicate that SHAPE reagents have similar half-life to 1M7 [38], and our results show that SHAPE-based methods align well with known RBP binding data (**Figure 6C-E**). RBP binding alters chemical probe reactivity in complex ways, either increasing or decreasing accessibility depending on the interaction [21]. These changes make it impossible for traditional methods, which rely on limited features, to distinguish RBP-bound regions from fully double-stranded RNA using single probing datasets (**Figure 6A**). By integrating 15 expanded features, STONE enables the differentiation of these intricate patterns, providing a novel approach for identifying RBP binding sites based solely on single probing experiments (**Figures 6**). Moreover, STONE can be further employed to non-targeted RBP binding sites identification and validated by molecular techniques such as pull-down assays, proteomics, and CLIP-seq. These molecular experiments enhance the reliability of the STONE-derived method and provide valuable insights for AI-based models of binding site identification [16].

Integrating both mutation and stop signals to improve RNA structure analysis accuracy represents a pivotal advancement, enabling not only the refinement of structural maps in existing datasets but also the expansion of potential applications. This improvement strengthens the foundation for numerous computational tools, such as PrismNet [20] and IntaRNA [39] which predict RNA interactions with biomolecules, and DiffScan [40], which assesses RNA structural heterogeneity across samples. By enhancing RNA structural signal accuracy, STONE directly impacts the reliability of these computational analyses and facilitates more robust downstream applications [9,41]. STONE integration method underscores the importance of precise structural insights for advancing RNA research and offers a platform for the continued development of AI-based models and experimental validations, bridging computational predictions with biological discovery.

### Limitations of the study

STONE, while presenting significant advancements in RNA structure analysis, also has certain limitations that must be addressed to fully realize its potential. First, the performance of STONE relies heavily on the integrity and balance of mutation and stop signals generated during RNA structure probing experiments. However, these signals can be variably affected by experimental workflows, particularly during library preparation steps that may selectively degrade stop signals. This dependency limits STONE’s applicability to datasets produced by specific probing strategies and highlights the need for methods that ensure consistent signal preservation across workflows.

Additionally, the compatibility of STONE with a broader range of chemical modification reagents remains an area for further exploration. While the method demonstrated robust performance with DMS and SHAPE reagents, its efficacy with other reagents, such as 2A3 or emerging chemical modifiers, has not been validated. Different reagents target distinct chemical environments within RNA, and variations in their half-lives, reactivity, and structural preferences could impact the integration of mutation and stop signals. Notably, the two models underlying STONE are structured to accommodate molecular differences, suggesting that they could be extended to other reagents. However, the lack of sufficient datasets prevents definitive evaluation at this stage. Expanding reagent compatibility is critical for extending STONE’s application to diverse biological and environmental conditions.

Another limitation is that current chemical probing methods, including those used by STONE, inherently lose information regarding long-distance RNA interactions. This limitation is a common issue with DMS- and SHAPE-based methods, which primarily capture local nucleotide pairing probabilities. Specialized techniques such as PARIS [42] and RIC-seq [43] have been developed specifically to capture long-distance interactions. However, the primary focus of our study is to enhance the accuracy of chemical probing methods rather than to capture long-range interactions; thus, the absence of long-distance interaction data is beyond the scope of this work.

Despite these limitations, STONE represents a transformative tool in RNA structural biology, with the potential to refine RNA structural maps and broaden the scope of computational RNA research. Future developments addressing these challenges will further enhance its utility and impact in the field.

## CONCLUSIONS

The effectiveness and complementarity of dual signals have been validated, laying the foundation for integrating these signals. The STONE method improves RNA structure analysis accuracy and enhances signal coverage both at the genome-wide and single-cell levels. STONE leverages small molecule accessibility insights to directly detect RBP binding sites from a single RNA structure probing experiment.

## MATERIALS & METHODS

### Data preparation

We downloaded raw data from publications included in the RASP database [35] and additional recent genome-wide RNA structure probing studies. The final datasets encompassed 11 studies, spanning five species, six small molecule reagents, and four reverse transcriptases (**Table S1**). For signal distribution calculation, all studies were included, excluding single-cell datasets. In evaluating well-characterized RNAs, we focused on human, mouse, virus, and yeast for structural analysis. For the genome-wide evaluation, we utilized high-depth representative datasets. The single-cell datasets included two batches of the H9 cell line, totaling 76 cells [15]. For the RBP binding site identification application, CLIP-seq data were compiled from our previous work [16], and RNP-MaP datasets were sourced from HEK293T cell line [33]. All data were downloaded to the computing platform server in the Core Facility and Service Platform, School of Life Sciences, Shandong University, with MD5 checksums performed to verify data integrity.

After obtaining the raw sequencing files in FASTQ format, low quality reads were removed using fastp software (version 0.32.4) with default parameters [44]. Reads were filtered using fastp with parameters set to remove duplicates (*--dedup*), applied a minimum Phred score of 20 (*--qualified_quality_phred=20*), exclude reads with >20% low-quality bases (*--cut_mean_quality=20*), and retain a minimum read length of 5 bases (*--length_required=5*). This rigorous preprocessing was essential for minimizing noise and increasing the reliability of subsequent analyses. Adapters were either auto-detected and removed or specified according to details in the original publications.

### Sequence alignment and signal calling

Bowtie2 (version 2.4) was used to build genome reference indexes (**Table S1**) and align reads to the reference genome [45]. The *--local* option was applied to optimize mapping efficiency. Resulting BAM files were then processed for mutation signal calling with RNA Framework (version 2.8.7) [34], with parameters configured to exclude insertions while retaining default settings for deletions. The *--orc* option was implemented to include raw mutation counts summarizing across mutation directions, insertions, and deletions. For stop signal calling, we followed the default settings recommended by the icSHAPE-pipe (version 2.0.2) [25]. Finally, we calculated stop and mutation rates based on count and depth at each position from the processed signals generated by icSHAPE-pipe and RNA Framework.

### Signal merging and processing

Genome-wide data processing requires three input files: two from RNA Framework analysis (*rf-count --orc*) and one from icSHAPE-pipe (*calcSHAPENoCont*). To facilitate downstream modeling, we developed command tools in Rust programming language to process and merge these files into the required format.

The output files generated by the *rf-count* command in RNA Framework were first processed into an intermediate format using the *zip_rfcsv* command. Raw counts files generated with *--orc* enabled were initially compressed with the *zip_rftxt* command, and then further processed into an intermediate format via the *zip_rftxt2* command. Similarly, output from the icSHAPE-pipe *calcSHAPENoCont* command was compressed into an intermediate format for stop signals with the *zip_pipe* command.

These intermediate formats were merged with the *merge* command, resulting in a comprehensive file where each row represented a specific RNA locus. This file recorded mutation count, sequencing depth, and mutation direction from RNA Framework, alongside stop signal counts from icSHAPE-pipe. Additionally, the *mbreport* command was applied to calculate the average depth and counts for both mutation and stop signals in genome-wide datasets.

To analyze well-characterized structures such as 18S rRNA, a streamlined approach was implemented using a custom R package named shapeTM (based on R version 4.3.2). In this workflow, the three output files described above were used directly as inputs without requiring the compression step, thus simplifying single-gene analysis.

### Feature engineering and model training

Based on the merged and processed signal data, we expanded the features to 42 types, as detailed in **Table S2**. After evaluating the learning capacity and generalization of the model across various feature combinations, we ultimately reduced the input to 15 features. For training the SHAPE model, we used 18S rRNA data from human [15,16,46,47], as well as mouse data [36]. Note that although the 2A3 dataset [46] was incorporated for training due to its high signal-to-noise ratio, it was not used for subsequent validation because fragment selection resulted in insufficient stop signals for effective dual-signal integration. For the DMS model, datasets included 18S rRNA data from *Arabidopsis thaliana* [3], human data [26], and yeast data [4,48]. Positive samples represented unpaired sites, while negative samples represented paired sites, yielding 7,106 positive and 6,907 negative samples for the SHAPE model, and 1,814 positive and 1,471 negative samples for the DMS model. The data were initially merged and subsequently partitioned into training and test sets, with 75% of the samples designated for training and the remaining 25% reserved for testing.

The training sets were applied to various machine learning algorithms, including Gradient Boosting, eXtreme Gradient Boosting, Decision Tree, and Random Forest, with hyperparameter tuning performed to optimize model performance. In terms of deep learning models, training was performed using CNN_ResNet50, CNN_Vgg16, CNN_Xception, Transformer, RNN, GRU, biGRU, LSTM, and biLSTM. After training, the models were evaluated on both the 25% test dataset and independent validation datasets. Model performance was assessed using several metrics, including AUC, F1 Score, Precision, and Accuracy. AUC values were calculated using the *roc_curve* and *auc* functions from the scikit-learn library (version 0.24.2). To further evaluate model generalization and potential overfitting, we employed 10-fold Cross-Validation, dividing the dataset into ten mutually exclusive subsets. For each training iteration, nine subsets were used for training, while the remaining subset was used as the test set. The optimal model was selected based on the highest AUC values achieved in both the validation and test datasets.

### STONE validation by known RNA structures

To mitigate the influence of outliers and ensure data comparability within a consistent range, structural scores output from both the traditional method and STONE models were normalized to a [0, 1] scale using quantile-based normalization [25]. Following the approach outlined by Zarringhalam et al. [49], a nonlinear transformation was applied to the normalized stop and mutation scores to adjust for weight differences across their respective numerical ranges.

Independent datasets were used to validate both the DMS and SHAPE models by comparing the structural scores with known RNA structures obtained from the RNA STRAND database (http://www.rnasoft.ca/strand, version 2.0) [50]. For DMS validation, datasets from DMS-seq [26], DMS-MaPseq [4], Mod-seq [48], and Structure-seq [3] were employed. For SHAPE validation, datasets from SHAPE-Seq [36], icSHAPE [16,51], SHAPE-MaP [52], and sc-SPORT [15] were used. Positions with missing structural scores were excluded, and the remaining positions were evaluated to determine whether they corresponded to unpaired nucleotides, allowing for the calculation of AUC values.

To evaluate RNA structure prediction using STONE scores, we employed RNAframework’s rf-jackknife method to optimize folding parameters [34]. Specifically, we performed a grid search to determine the optimal slope and intercept values for both the STONE score and the original signals. These optimized parameters were then used to fold the RNA by Superfold (version 1.0) [6], and the predicted structures were compared to reference structures obtained from the RNA STRAND database.

The resulting structural scores (stop, mutation, and STONE) were visualized in the Integrative Genomics Viewer (IGV version 2.16.2) [53], along with the reference sequence, to display structural signals at single-nucleotide resolution. A custom Python script was used to map these scores onto the secondary RNA structure of human 18S rRNA, facilitating a visual evaluation of scoring accuracy across methods.

Accessibility information was derived from the 60S ribosome crystal structure (PDB ID: 5LKS), and the solvent-accessible surface area for each nucleotide was calculated using PyMOL (version 3.0.3).

### Genome-wide analysis

For genome-wide analysis, high-depth representative H9 datasets from Sun et al. [16] were selected. Gene longest transcript was extracted from the human reference genome (GCF_000001405.40, GRCh38.p14) for subsequent signal quantification. To assess model performance on human 18S rRNA across datasets with varying depths, the SAMtools (version 1.15.1) *view* command was used to subset mapped reads in the BAM file at designated percentages (0.5, 0.25, 0.1, 0.05, 0.025, 0.01, 0.005, 0.0025, 0.001, 0.0005, 0.00025, 0.0001, and 0.00005). After signal calling, the average depth of all nucleotides was calculated and plotted against the corresponding AUC values. To quantify gene number improvement with reliable RNA structure signals in the H9 dataset conducted by icSHAPE [16]. The AUC value before downsampling was used as baseline. Effective coverage was defined in two aspects—gene-level and nucleotide-level. Specifically, we established the minimum read depth threshold as the depth at which the STONE AUC declines to 80% of its maximum value observed in the undownsampled dataset. Gene-level coverage was then determined by counting the number of genes with average read depths meeting or exceeding this threshold, while nucleotide-level coverage was calculated by counting the number of transcript positions with non-empty STONE and original stop signals (**Figure S11**). For example, the STONE method achieves an AUC of 0.755 at the highest depth, and the effective coverage reaches this 80% threshold (AUC = 0.604) at 35 reads. At this depth, the nucleotide coverage is 1.8 × 10. In comparison, for the original method to reach the same AUC accuracy, the depth required is 150 reads, corresponding to a nucleotide coverage of 6.8 × 10.

Based on the downsampling results, which demonstrated STONE’s ability to improve accuracy in low-depth datasets, we further analyzed its potential for genome-wide RNA structure studies. This included examining RNA structural features and associated transcript-level information, such as gene transcription levels and RBP-target RNA interactions. Initially, the mapping output SAM files were converted to BAM format using SAMtools (version 1.15.1). The BAM files were then sorted by genomic coordinates to ensure accuracy and efficiency in subsequent Fragments Per Kilobase of transcript per Million mapped reads (FPKM) calculations. FPKM values for each transcript were computed using Cufflinks (version 2.2.1) [54]. From the FPKM results, transcripts with values greater than 0 were extracted, sorted in descending order, and saved in CSV format for further analysis. Then, the intensity of STONE, mutation, and stop scores for each transcript was visualized in three independent lanes of circular plot, sorted in descending order based on their FPKM values, using a template generated by itol.toolkit (version 1.1.8) [55] in iTOL (version 6.9.1) [56]. Relationships between RBPs and their target RNAs were calculated using LIN28A and TARDBP datasets [57] from the H9 cell line and visualized in the circular plot.

The structure motif search focused on stem loops with stem lengths ranging from 2 to 7 nucleotides (nt) and loop lengths between 2 and 7 nt. RNA structural scores and the corresponding transcripts sequences were used as input for structural motif identification. To analyze the presence of structural motifs, PatteRNA (version 2.1) was employed [31]. Only stem-loop motifs with a positive probability, indicating a higher likelihood of actual occurrence based on the structural score, were considered for further analysis.

### Single-cell analysis

The single-cell dataset has two batches of the H9 cell line totaling 76 cells available [15]. The structural scores were also calculated using the longest transcripts as references. The method for calculating RNA structural heterogeneity was as described in the same study [15]. The number of genes and nucleotide site coverage were determined as described previously in the genome-wide analysis method section.

To explore the characteristics of real resampling in single cells and to compare the dependence of different methods on data depth, an NLS model was employed to fit the average sequencing depth and covered nucleotides percentage for the same transcript through single-cell multiple sampling analysis.

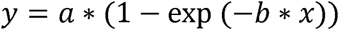

Here, y represents the percentage of covered nucleotide sites (dependent variable), while x is the average sequencing depth of the transcript (independent variable). The parameter a denotes the asymptote, representing the maximum of covered nucleotides percentage as x increases. The parameter b controls the rate at which the coverage approaches the plateau, with higher values of b indicating faster convergence.

One H9 cell dataset (sample id RHS3588) was used as a representative sample for RNA structure folding. Meanwhile, a genome-wide bulk dataset from the H9 cell line [16] was also folded to generate a reference structure to ensure structure reliability. STONE and original scores were incorporated as constraints to refine RNA folding using the Minimum Free Energy (MFE) approach, implemented in Superfold (version 1.0) [6]. During the Superfold folding process, the partition function for each available transcript was calculated automatically to formulate arc plots. Pairing similarity between the single-cell and bulk structures was quantified by calculating the number of common base pairs from arc plots. The matching ratio was then defined as the proportion of matched base pairs relative to the total base pairs in the bulk structure, providing a measure of structural similarity.

### RBP binding site identification

To calculate ΔSTONE values, the STONE output was compared with original signals, strictly following the previously established method [21,24]. Positions exhibiting different signals were identified (2964 positions) using a sliding window approach, and consecutive positions were grouped into regions (628 regions). Specifically, within each window (size=5 nt), ΔSTONE-identified positions were considered valid only when at least three positions were identified. The steps for calculating ΔSTONE are outlined below.

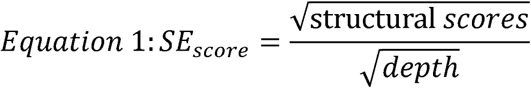

In **Equation 1**, **depth** refers to the read depth at a specific nucleotide, and **structural scores** represent the STONE score, mutation score, or stop score. The standard errors (**SE_score_**) for each score were calculated.

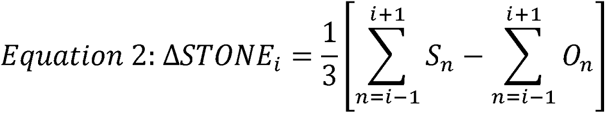

The structural score difference (Δ**STONE**) for each nucleotide was determined using **Equation 2**, where ***S*** and ***O*** represent the structural scores obtained from STONE and original methods, respectively. The ΔSTONE values captured the difference in scores and were averaged across a three-nucleotide sliding window.

To incorporate smoothing, the standard error values obtained from **Equation 1** were averaged, ensuring alignment with the methodology established by Smola et al. [24]. The smoothed error estimation for each nucleotide position was denoted as *SE_i_*.

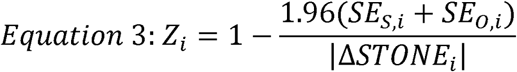

In **Equation 3**, *Z*-Factors (*Z*) for each nucleotide (***Z_i_***) were calculated, where the subscripts ***S*** and ***O*** represent the STONE and original method (according to **Equation 2**), respectively. Nucleotides with *Z* > 0 were considered to exhibit significant changes in SHAPE reactivity.

Standard scores (**S_i_**) for each nucleotide were rigorously computed, and the identification of potential binding sites was performed strictly according to the methodology outlined by Smola et al. [24].

### Evaluation of **Δ**STONE results

The CLIP-seq data were obtained from the POSTAR3 CLIPdb database in BED format (http://111.198.139.65/RBP.html, last accessed on Nov 15 2024) [57]. Six experimental methods were utilized: PAR-CLIP, PIP-seq, eCLIP, HITS-CLIP, iCLIP, and 4SU-iCLIP. To extract data for specific genes, the CLIP-seq RBP binding sites were annotated to the corresponding transcriptome using GAP (https://github.com/lipan6461188/GAP, version 1.0).

RNP-MaP sites extraction was based on the methodology described by Weidmann et al. [21]. The processed data files (GEO accession ID GSE152483) include mutation frequencies and read depths for both crosslinked and uncrosslinked samples. All computations were implemented in a custom R script and generated candidate RNP-MaP sites on human XIST lncRNA.

To validate the accuracy of ΔSTONE calculations, ΔSTONE outcomes were compared with CLIP-seq and RNP-MaP results. In this analysis, consecutive nucleotide positions within ΔSTONE, CLIP-seq, and RNP-MaP data were grouped to independently represent their RBP binding regions. To minimize the likelihood of random overlap, two criteria were applied when determining overlapping regions: for CLIP regions shorter than or equal to 6 nt, an overlap was considered valid if the overlapped length exceeded 3 nt; for CLIP regions longer than 6 nt, the overlap threshold was set to a minimum of 5 nt to be considered valid.

The statistical significance of the observed overlaps was assessed by generating randomly sampled nucleotides. Specifically, 2,964 nucleotides were randomly selected, equivalent to the number of ΔSTONE-identified binding positions. For each RBP, ten sets of randomly sampled positions matching the number of CLIP-identified binding positions were generated to examine the statistical significance by one sample t-test. For visualization, the midpoints of the CLIP-identified regions were used to represent each region in the plots, providing a clear representation of the spatial distribution of the RBP binding areas.

## QUANTIFICATION AND STATISTICAL ANALYSIS

All statistical analyses were performed with R. Where applicable, the sample size (n) was specified either in the plot or in the corresponding figure legends. Analysis details could be found in the method section, such as statistical tests, definitions, etc. Boxplots were used to visualize the distribution of the data, with the box representing the interquartile range (IQR). The median was indicated by a horizontal line within the box, while the upper and lower hinges represented the 75th and 25th percentiles, respectively. The whiskers extended to the most extreme data points within 1.5 times the IQR from the box. Any data points beyond this range were considered outliers and plotted individually. Statistical significance was determined using appropriate tests, with p-values reported in the figure legends or main text.

---

**Supplementary Figure 1.**
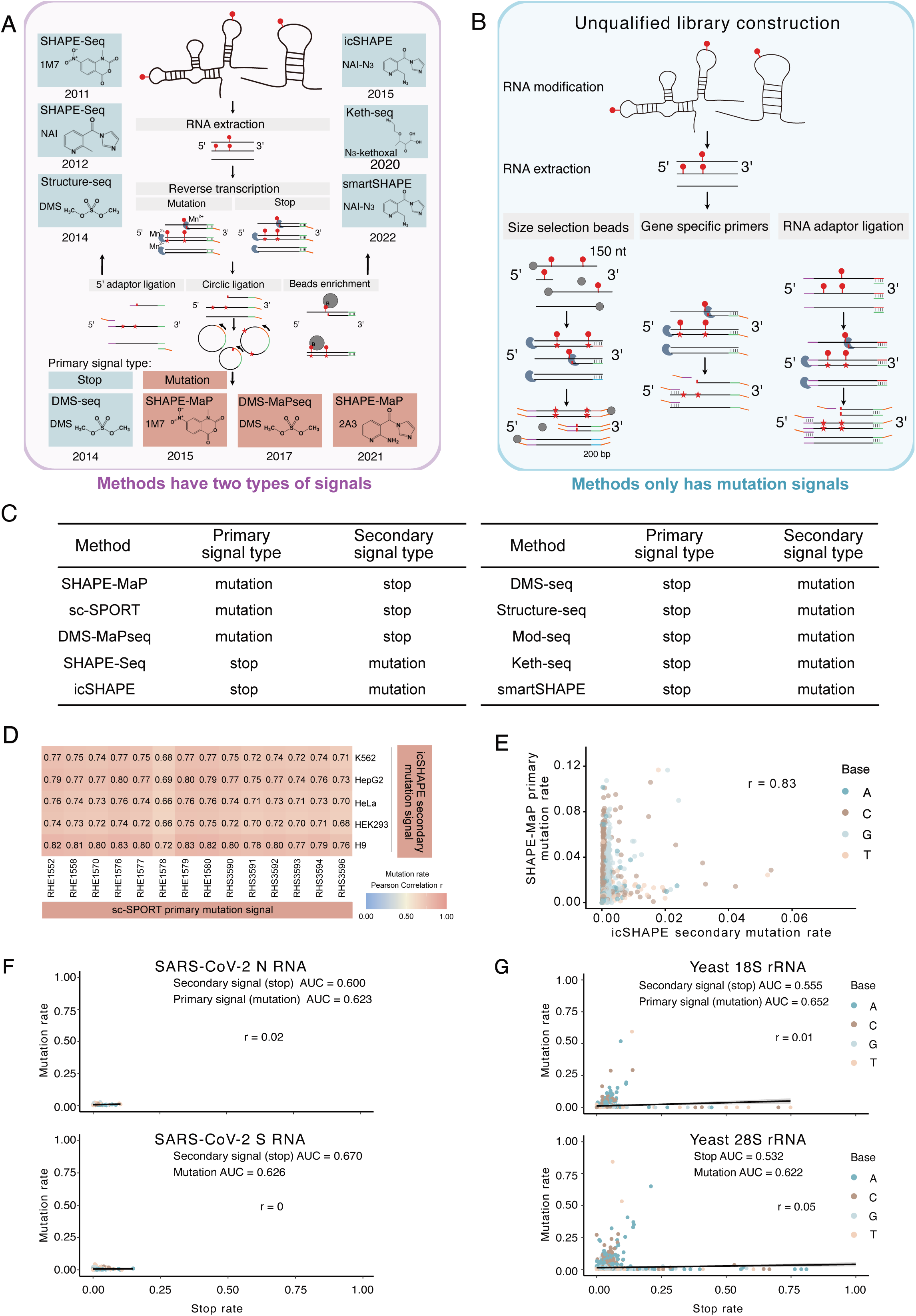
Signal distributions of primary and secondary signals within one experiment. (A) Overview of commonly used RNA structure probing workflows. Experimental methods are color-coded by their original experimental strategies: cyan for stop-based methods and orange for mutation-based approaches. Small molecule modification sites are indicated by red dots. (B) Library construction methodologies yielding only one type of signal, primarily stop signals. (C) Primary and secondary signal types of chemical probing methods. (D-E) The Pearson correlations of mutation signals for human 18S rRNA were calculated using results from two experimental strategies: sc-SPORT, where mutation signals were considered as primary signals, and icSHAPE, where mutation signals were treated as secondary signals. (F-G) Spearman correlation analysis of primary mutation and secondary stop signals was performed for yeast 18S and 28S rRNA in DMS-MaPseq dataset [4] and virus N and S RNA in SHAPE-MaP dataset [52]. The corresponded accuracies for both types of signals were independently calculated and annotated on each plot. Statistical significance is denoted as follows: n.s. (not significant) for p > 0.05, * for p < 0.05, ** for p < 0.01, *** for p < 0.001, **** for p < 0.0001.

**Supplementary Figure 2.**
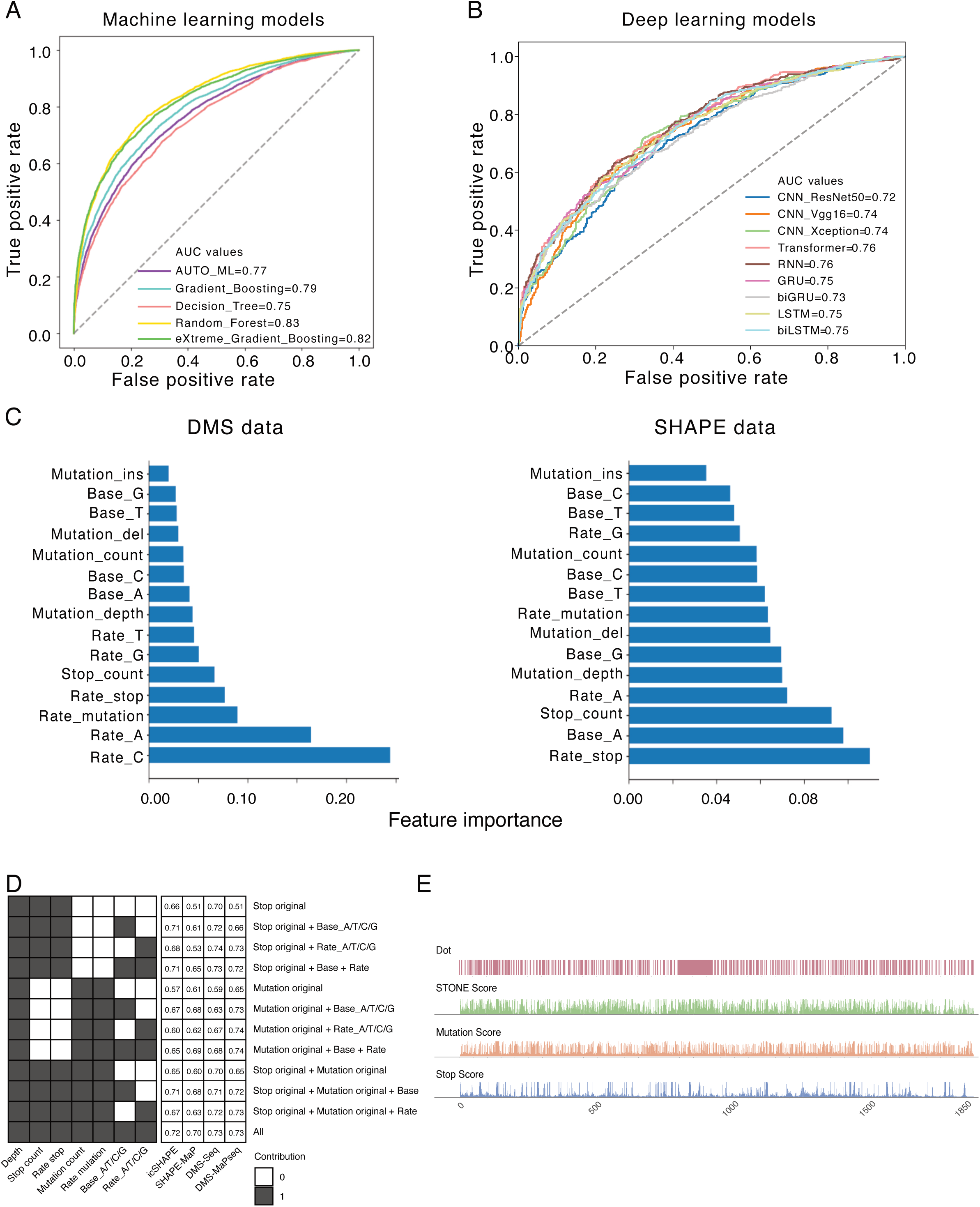
Additional background information and preliminary model outcomes. (A) AUC comparison across multiple machine learning models, utilizing 18S rRNA datasets to assess model performance. (B) AUC comparison across multiple deep learning models, utilizing 18S rRNA datasets to assess model performance. (C) Feature importance applied in the Random Forest model when processing DMS and SHAPE datasets. (D) Quantification of feature contributions towards final STONE AUC values using four datasets, including DMS-seq [26], DMS-MaPseq [4], icSHAPE [16], and SHAPE-MaP [15]. (E) Comparative analysis of structural scores derived from the original and STONE methods applied to the H9 cell line [16], visualized relative to the established 18S rRNA structure using IGV.

**Supplementary Figure 3.**
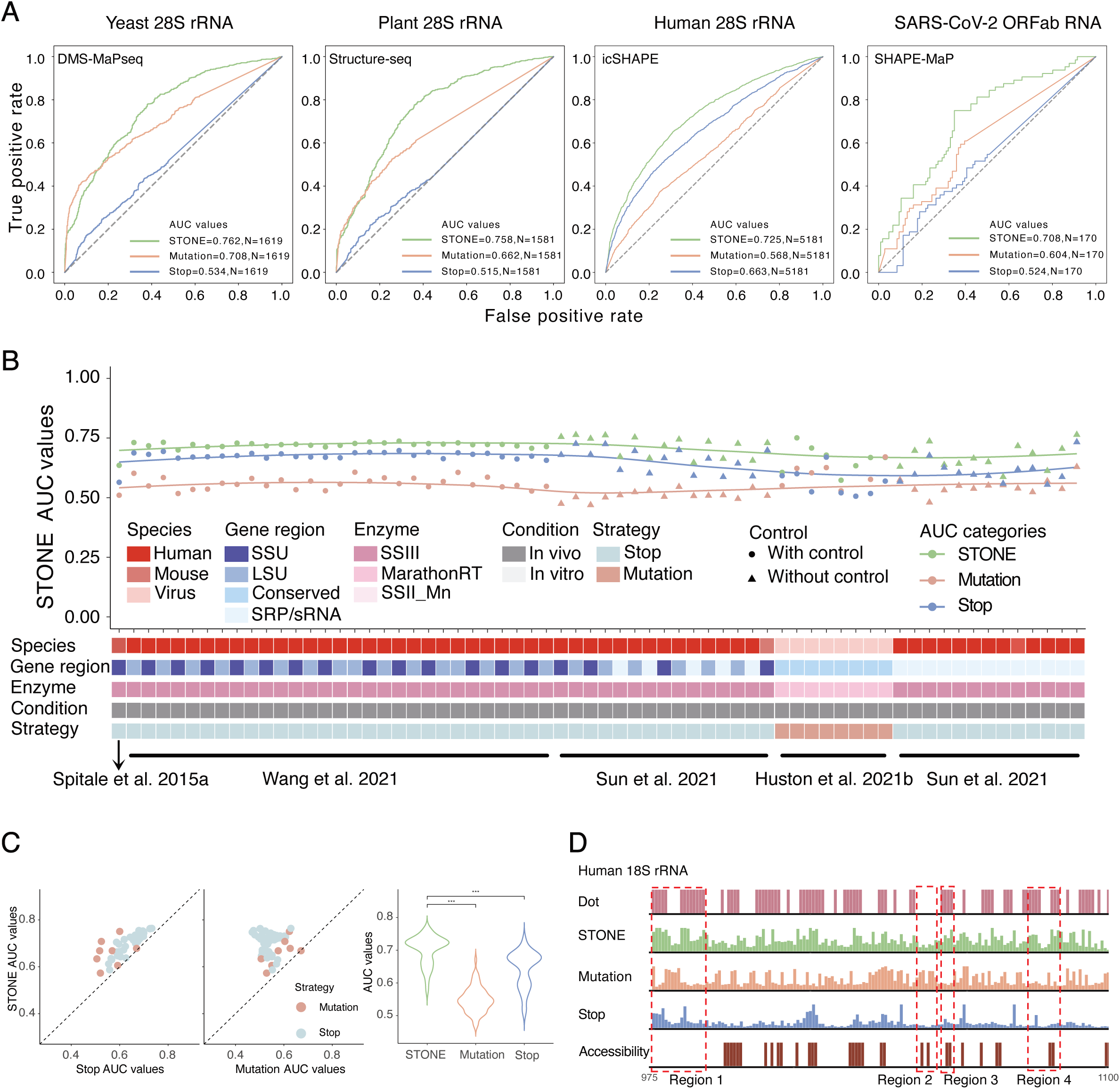
Large-scale validations on STONE AUC calculations. (A) Comparison of AUC values for original and STONE calculations applied to yeast 28S rRNA, human 28S rRNA, plant 28S rRNA, and SARS-CoV-2 open reading frame ab (ORFab) RNA. (B) Extensive validation of SHAPE model performance via AUC values across multiple datasets, including various species, gene regions, small molecule-enzyme combinations, experimental conditions, and reverse transcription strategies. (C) Statistical analysis of AUC values from 66 datasets, providing a comprehensive evaluation of the SHAPE model across various experimental contexts. The p-values were calculated using paired t-tests (n=66). (D) Single-nucleotide resolution improvements in RNA structural scores for human 18S rRNA. Results were evaluated using IGV. The red boxes highlighted successive positions with evident improvements.

**Supplementary Figure 4.**
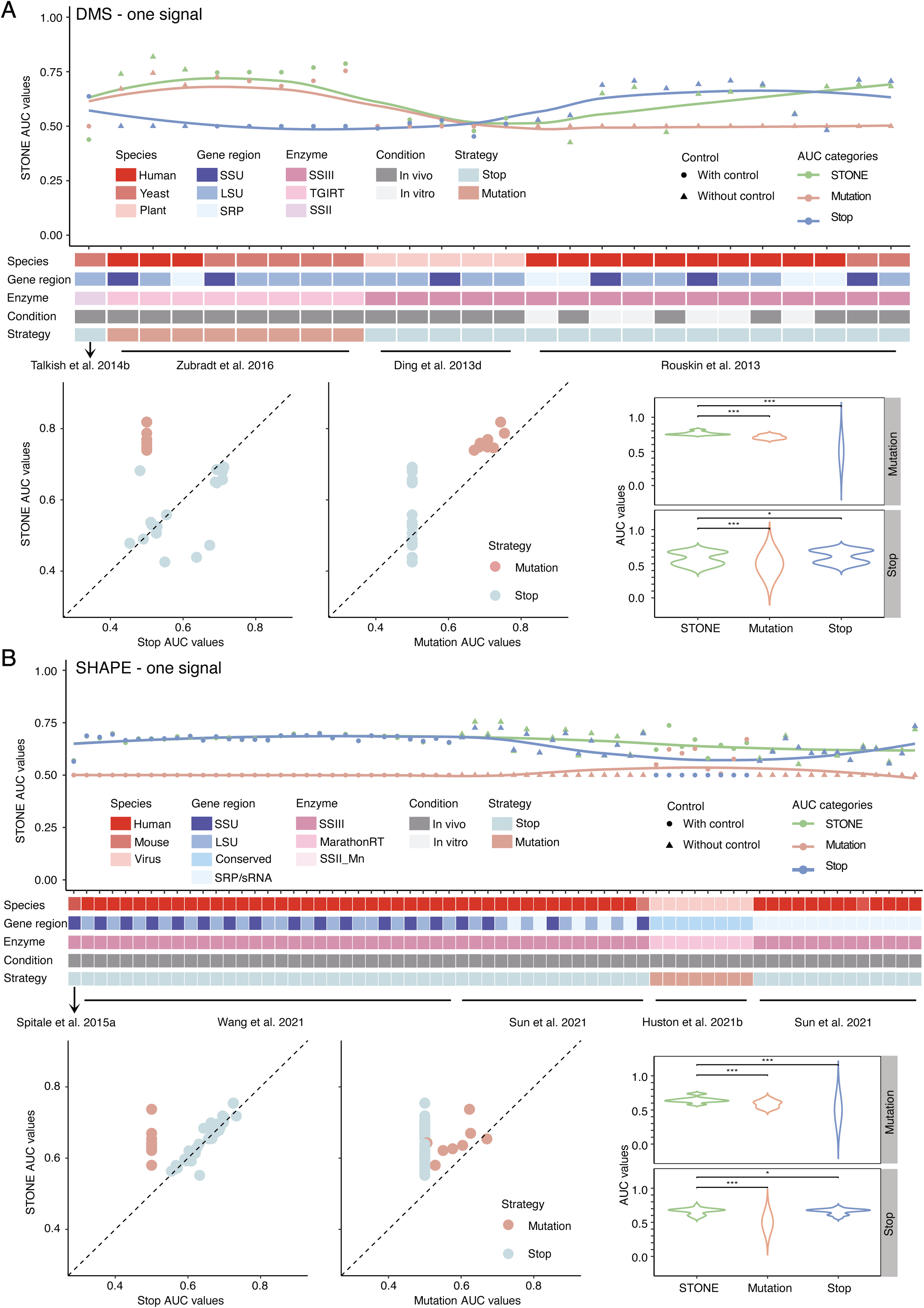
Examination for the potential overfitting issue by using one of the two signals. (A) Mitigation of STONE (DMS) overfitting risks by evaluating performance using one type of signal. The p-values were calculated using paired t-tests (n=26). (B) Mitigation of STONE (SHAPE) overfitting risks by evaluating performance using one type of signal. The p-values were calculated using paired t-tests (n=66).

**Supplementary Figure 5.**
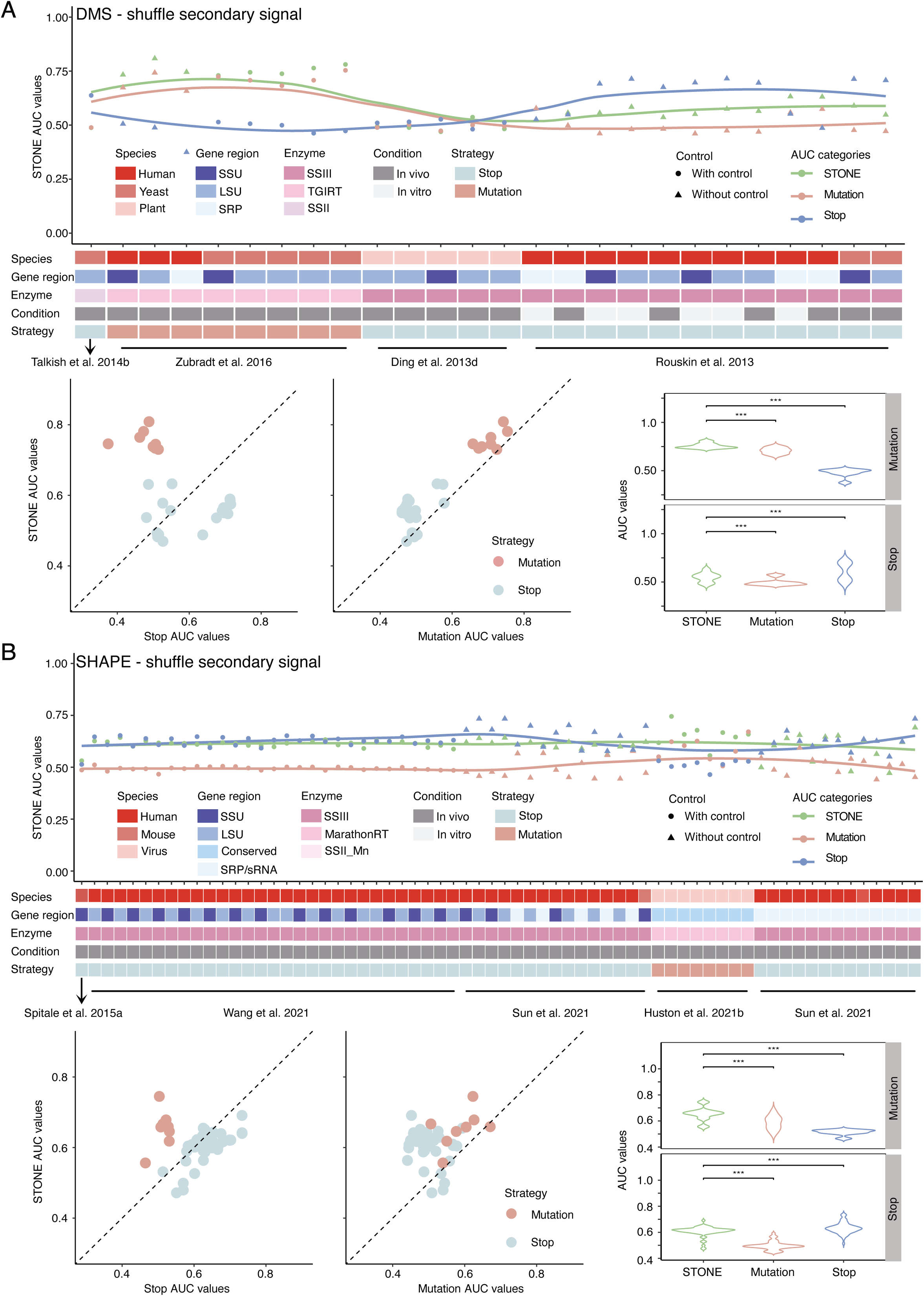
Examination for the potential overfitting issue by shuffling secondary signals. (A) Mitigation of STONE (DMS) overfitting risks by evaluating performance using shuffled secondary signal (n=26). (B) Mitigation of STONE (SHAPE) overfitting risks by evaluating performance using shuffled secondary signal (n=66).

**Supplementary Figure 6.**
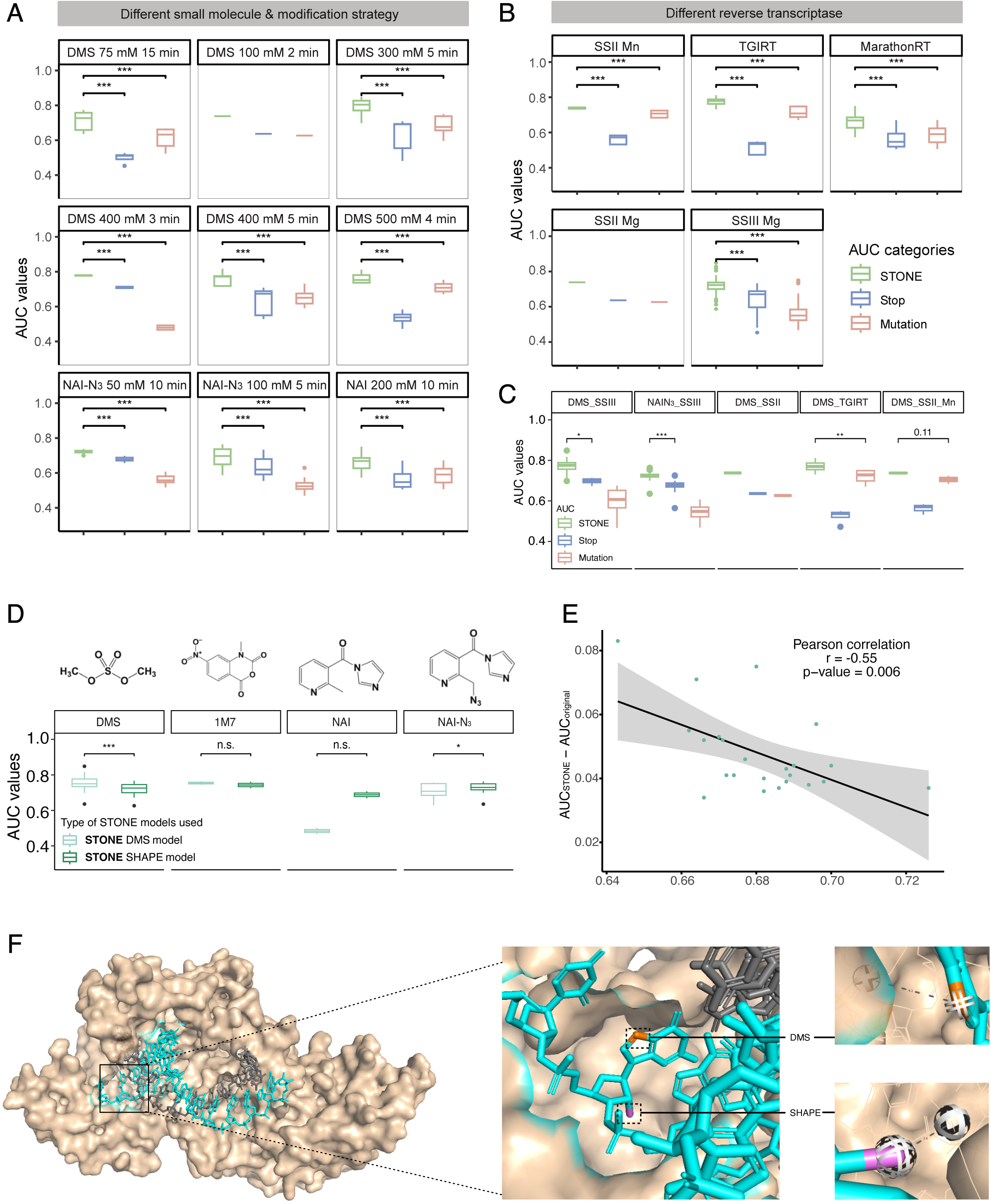
STONE performance for calculating human 18S and 28S rRNAs structures across different experimental protocols with varying small molecule modification conditions and reverse transcriptases. (A) AUC values under different small molecule modification strategies, including variations in reagent concentration, reaction time, and modification type, were calculated. AUC categories are color-coded: STONE (green), Stop (blue), and Mutation (red). Statistical significance was assessed using paired t-tests, with corresponding p-values displayed in the plots. (B) AUC values obtained from results produced by different reverse transcriptases, demonstrating consistency in model performance across various enzymatic conditions. Statistical significance was assessed using paired t-tests, with corresponding p-values displayed in the plots. (C) Evaluation of model performance across different combinations of small molecules and reverse transcriptases adopted for RNA structure probing experiments. Paired t-tests were employed to assess the significance of differences in AUC values. The number of datasets used for each combination is as follows: n=8 for DMS_SSIII, n=40 for NAI-N_3__SSIII, n=1 for DMS_SSII, n=4 for DMS_TGIRT, and n=3 for DMS_SSII_Mn. (D) Comparison of performance between DMS and SHAPE models in response to different small molecule modifications. The chemical structures of the small molecules are also displayed. The p-values were calculated using paired t-tests. The number of datasets used for each modification is as follows: n=2 for 1M7, n=3 for NAI, n=19 for DMS, and n=14 for NAI-N_3_. (E) Correlation between the intensity of AUC improvements using the STONE method and the AUC values calculated using the original analysis pipeline. The analysis was based on datasets containing dual signals. The p-value is derived from Pearson correlation calculation. (F) Visualization of RNA-reverse transcriptase interactions. Structures were obtained from PDB (ID: 4OL8) and visualized using PyMOL. Reverse transcriptase is shown in wheat, RNA in blue, DNA in gray, the adduct site for DMS modification in orange, and the adduct site for SHAPE modification in purple. The minimum distances from adduct sites to the reverse transcriptase surface were annotated.

**Supplementary Figure 7.**
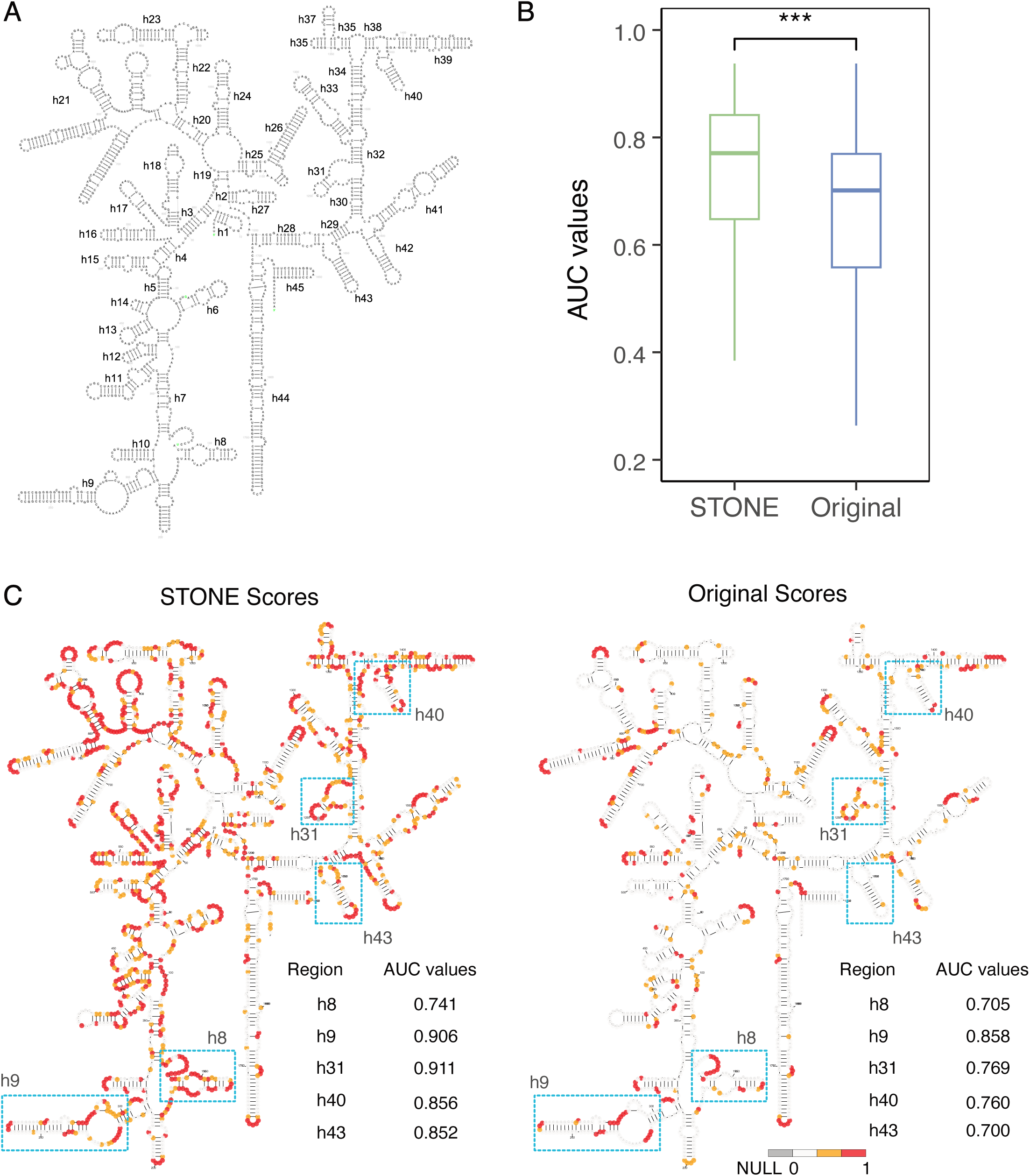
Regional STONE AUC improvement in human 18S rRNA. (A) Diagram illustrating the dissection of human 18S rRNA into 45 structural unit regions. (B) AUC values for the 45 regions of human 18S rRNA, calculated using STONE and traditional analysis pipeline. The dataset used was published by Sun et al. Statistical significance was assessed using paired t-tests, with corresponding p-values displayed in the plots. (C) Visualization of the accuracy and coverage of structural scores derived from the original and STONE methods published by Sun et al. [16] on the established 18S rRNA dot-bracket structure. AUC values for regions highlighted by blue boxes were independently calculated and annotated on the plot. Nucleotides were colored using structural scores.

**Supplementary Figure 8.**
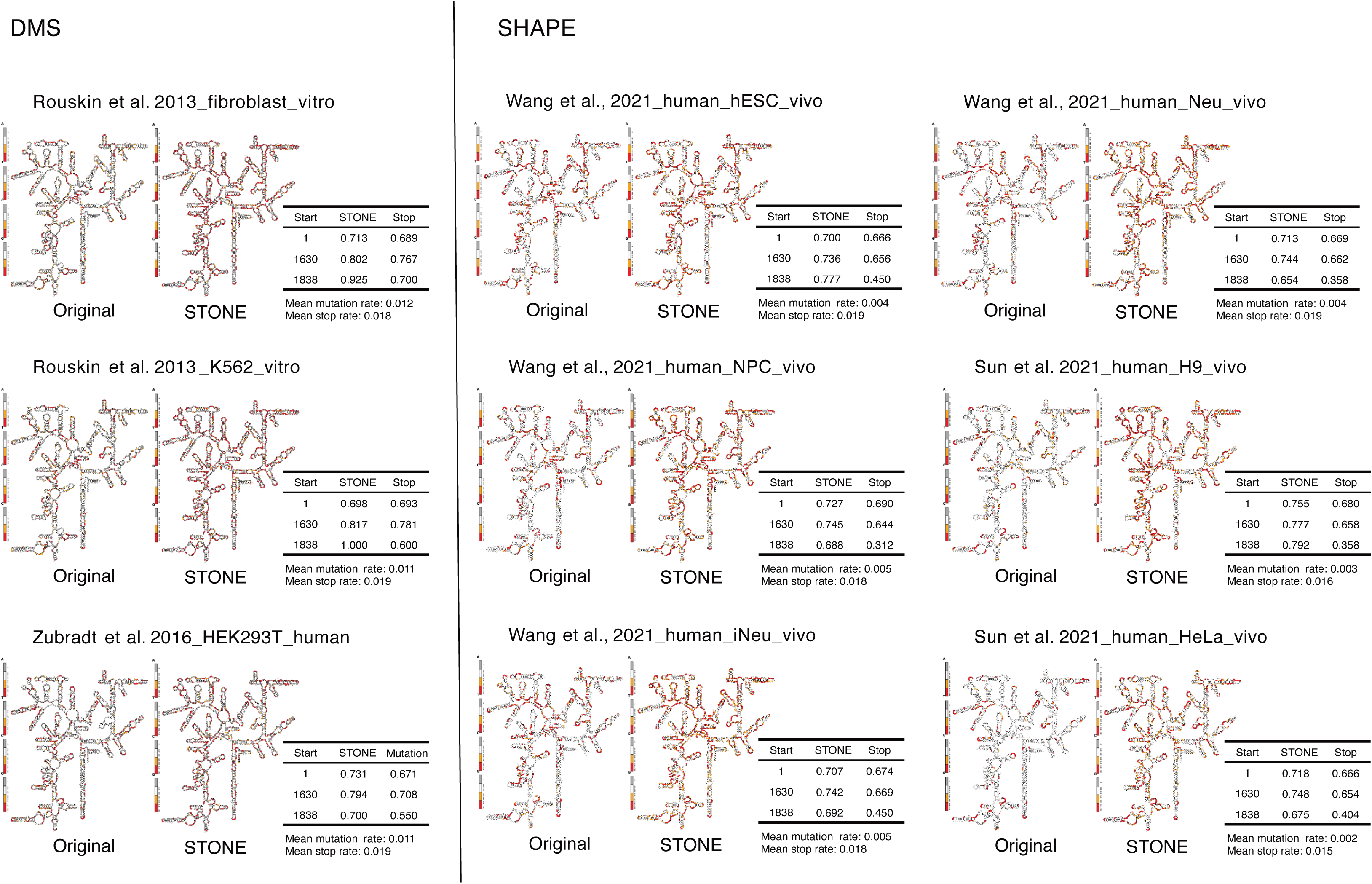
Visualization of structural scores from various datasets on established human 18S rRNA structure. Nucleotides were colored using structural scores.

**Supplementary Figure 9.**
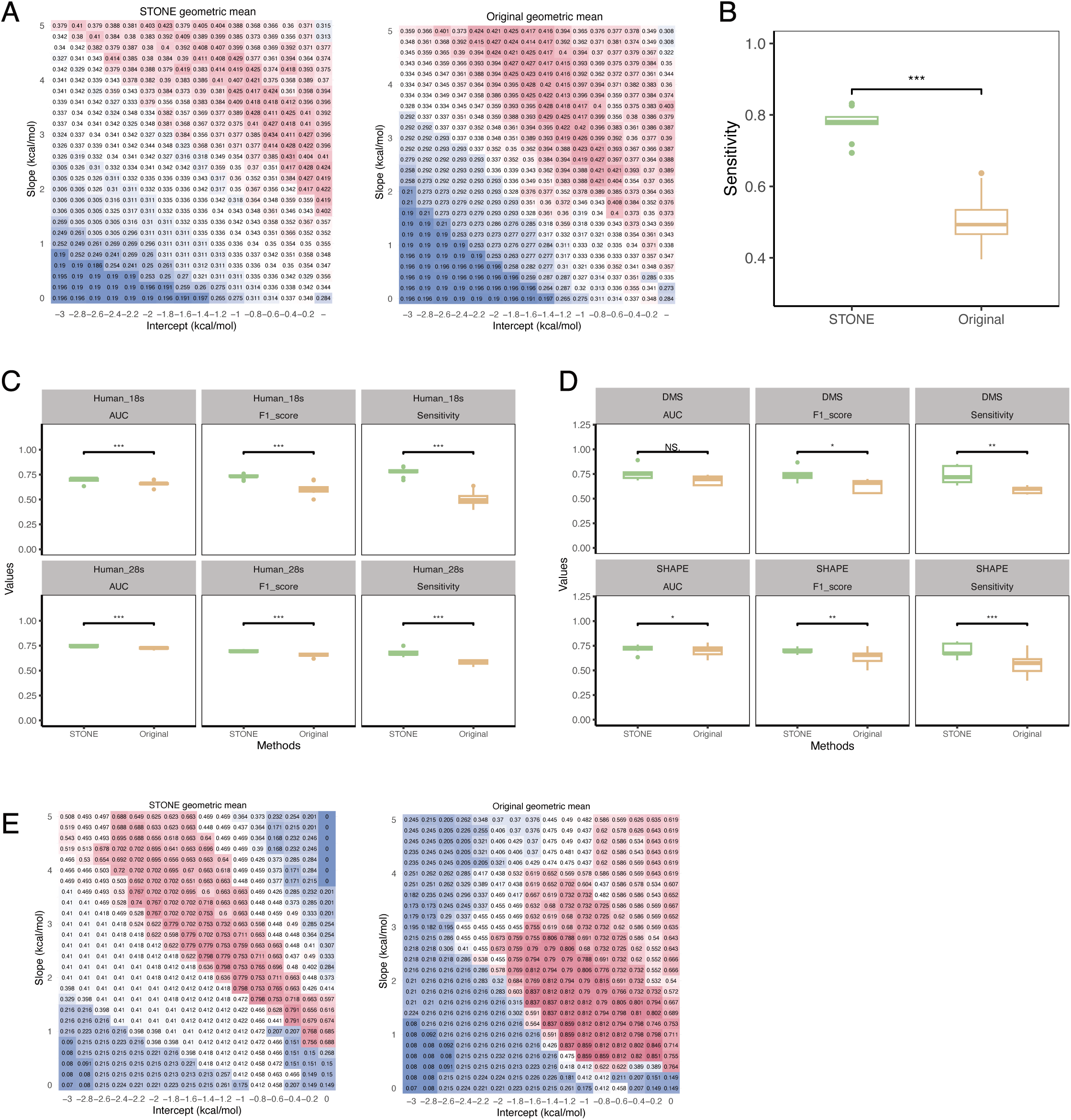
Structural prediction accuracy evaluation using STONE-derived structure scores. (A) Grid search for the optimal folding parameters for human 18S rRNA using STONE and the original structure scores. (B) Comparison of structural predictions for human 18S rRNA using optimized folding parameters derived from 10 datasets, including DMS-seq, DMS-MaPseq, SHAPE-Seq, and SHAPE-MaP. Sensitivity scores were calculated by comparing the STONE-derived and original method-derived structures with the known reference structure. Statistical significance was assessed using paired t-tests, with corresponding p-values displayed in the plots. (C) Evaluation of the accuracy of structural predictions for human 18S and 28S rRNA under various performance metrics. The box plots were derived from 22 datasets, including DMS-seq, DMS-MaPseq, icSHAPE, and SHAPE-MaP data. Predicted structures obtained using STONE analysis and those derived from traditional methods were compared to known structures. Statistical significance was assessed using paired t-tests, with corresponding p-values displayed in the plots. (D) Evaluation of the accuracy of structural predictions for DMS and SHAPE datasets under various performance metrics. The box plots were derived from 30 human datasets, containing various experimental methods (DMS-seq, DMS-MaPseq, icSHAPE, SHAPE-MaP) and distinct gene regions (18S, 28S, SRP, U1). Predicted structures obtained using STONE analysis and those derived from traditional methods were compared to known structures. Statistical significance was assessed using paired t-tests, with corresponding p-values displayed in the plots. (E) Grid search for the optimal folding parameters for human SRP RNA using STONE and the original structure scores.

**Supplementary Figure 10.**
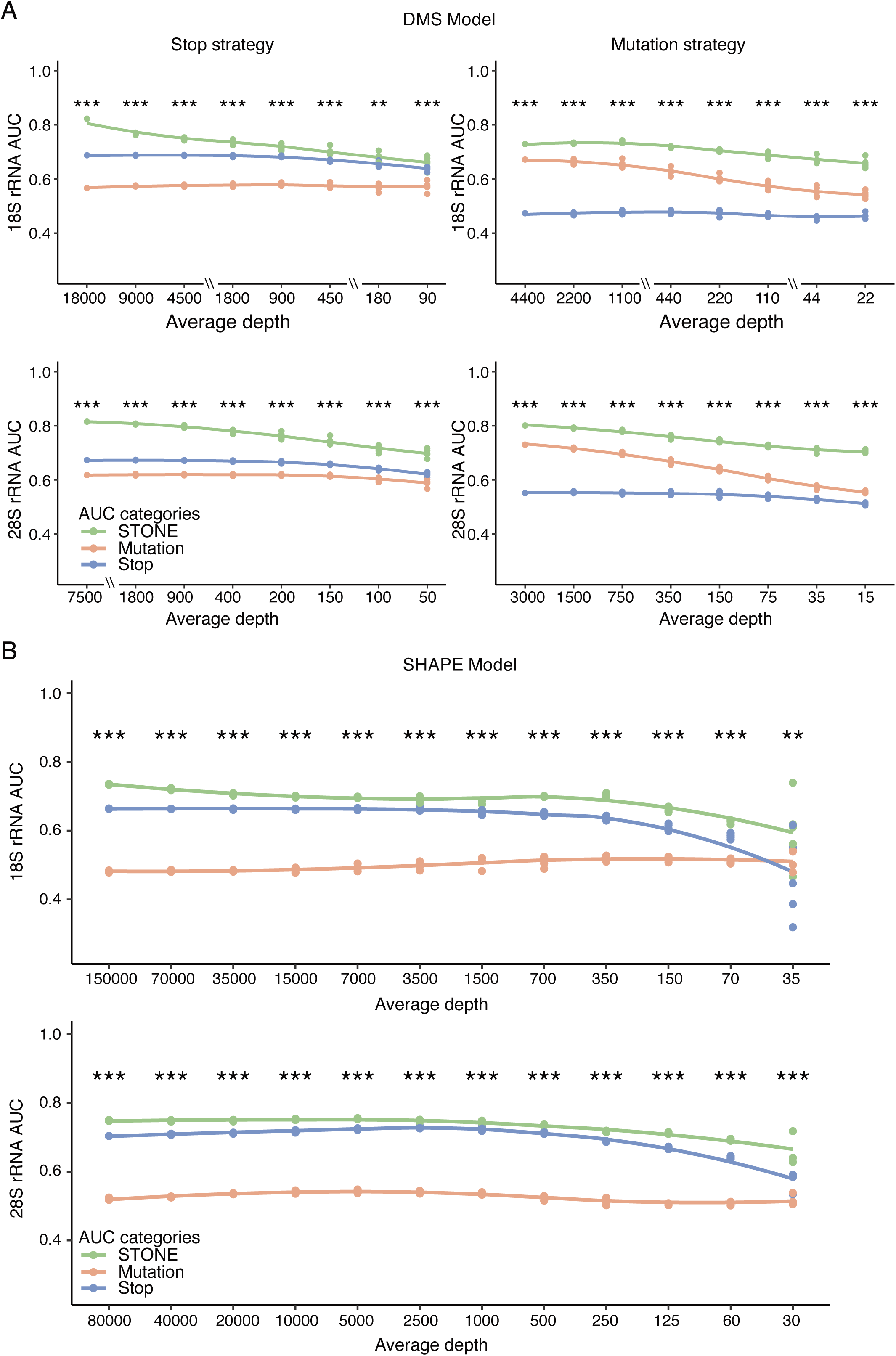
Additional down-sampled datasets for model robustness evaluation. (A) DMS datasets adopting stop (Rouskin et al. [26]) or mutation strategies (Zubradt et al. [4]) were downsampled for testing DMS model robustness on human 18S and 28S rRNA. (B) The SHAPE dataset adopting the stop strategy (Sun et al. [16]) was downsampled for testing SHAPE model robustness on human 18S and 28S rRNA. The p-values were calculated using paired t-tests to assess the significance of differences between STONE and original calculations (n=5). Statistical significance is denoted as follows: n.s. (not significant) for p > 0.05, * for p < 0.05, ** for p < 0.01, *** for p < 0.001.

**Supplementary Figure 11.**
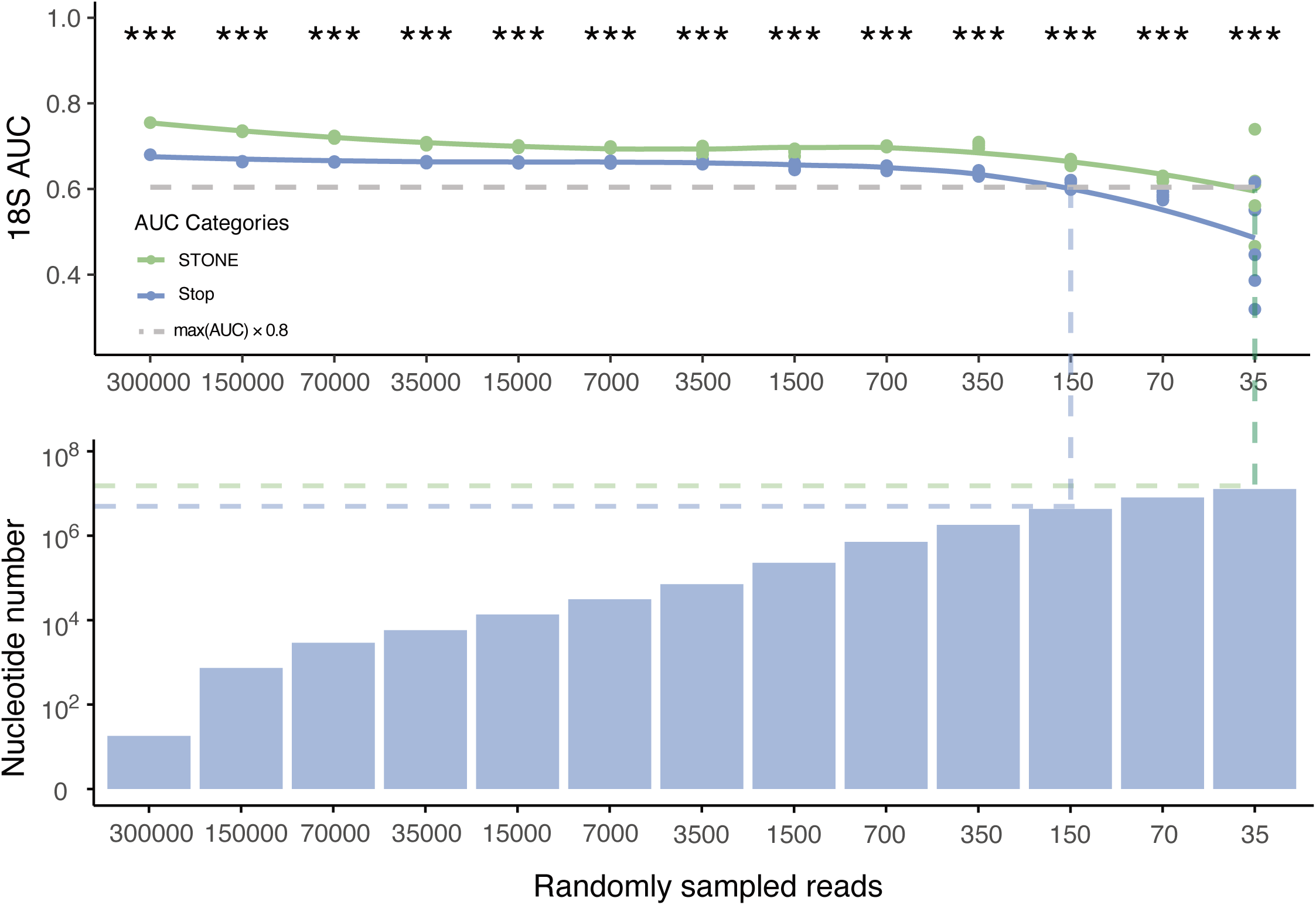
Nucleotide coverage obtained using STONE and original analysis. Nucleotide coverage refered to the number of nucleotides with effective signals at the minimal sequencing depth. The minimal sequencing depth for obtaining effective signals was defined as the depth at which the AUC falls to 80% of its maximum value under the highest sequencing depth (grey dashed line). For example, the STONE method achieves an AUC of 0.755 at the highest depth (300,000 reads), and the effective coverage reaches the 80% threshold (AUC = 0.604) at sequencing depth of 35 reads, corresponding to a nucleotide coverage of 1.8 × 10 (green dashed line). In comparison, the original method requires a minimum sequencing depth of 150 reads to achieve the same AUC accuracy, resulting in a nucleotide coverage of 6.8 × 10 (blue dashed line).

**Supplementary Figure 12.**
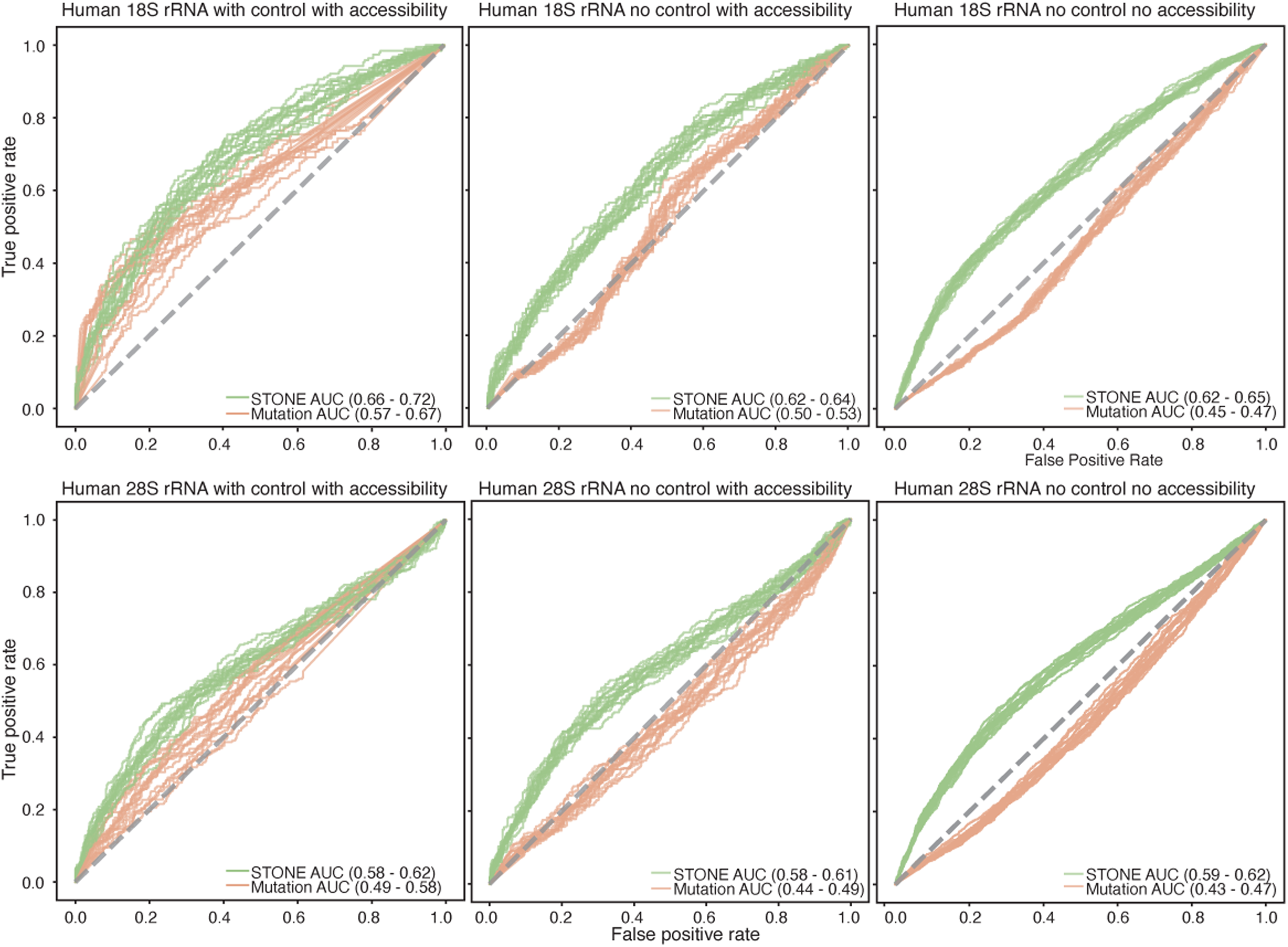
Comparison of STONE and original method performance on single-cell data. AUC values for human 18S and 28S rRNA calculated when considering different information combinations (n=15).

## TABLE LEGENDS

**Supplementary Table 1. Metadata of the publications and datasets considered and used in this study.**

**Supplementary Table 2. Description of features used in STONE calculation.**

## Acknowledgements

We thank Jindong Sun, Yongkang Tang and Chengqian Wang for fruitful discussions and invaluable feedback. We thank Ziyou Luo, Suiru Lu and Jingwen Ding for running the software comparison benchmark. We also thank Kui Xu, Jinsong Zhang, Wenze Huang, Jiale He, Yuhua Jiao and Yilin Song for critically reading the manuscript.

## Author’s Contribution

T.Z., R.Z., and S.Y. contributed equally as co-first authors and have agreed that either author can be listed first in reporting this study. The project was conceptualized by L.S. T.Z. established the initial technical framework and coordinated the specific task assignments for project progression. T.Z., R.Z., and S.Y. performed the computational analyses, data curation, and methods development. R.Z. trained the machine learning models, Q.C. trained the deep learning models, while S.Y. collected and re-analyzed the RNA structurome data. Y.H. developed high-performance computing tools in Rust. J.W. collected and re-analyzed RBP binding site data based on CLIP-seq, and X.T. performed the collection and re-analysis of RBP binding site data from RNP-MaP. Y.W. conducted key literature reviews. S.Y., Y.H., and Y.W. contributed to the chemical and structural biology visualizations. The initial draft of the manuscript was written by L.S., T.Z., S.Y., and R.Z., with C.L., L.S., S.Y., and Y.W. providing critical feedback and suggestions for improvement. S.Y. formatted the figures, and the final version of the manuscript was prepared by T.Z. and L.S.

## Funding

This work was supported by the National Natural Science Foundation of China (No.82341086, No.32300521, and No.32422013 to L.S.); the Open Grant from the Pingyuan Laboratory (No.2023PY-OP-0104 to L.S.); the State Key Laboratory of Microbial Technology Open Projects Fund (No.M2023-20 to L.S.); the Intramural Joint Program Fund of the State Key Laboratory of Microbial Technology (NO.SKLMTIJP-2024-02 to L.S., C.L., and T.Z.); the Shandong Province Postdoctoral Innovation Project (NO. SDCX-ZG-202400146 to T.Z.), the Qingdao Postdoctoral Science Foundation (NO. QDBSH20240102200 to T.Z.), the Double-First Class Initiative of Shandong University School of Life Sciences; the Young Innovation Team of Shandong Higher Education Institutions, the Taishan Scholars Youth Expert Program of Shandong Province, and the Program of Shandong University Qilu Young Scholars.

## Availability of data and materials

An implementation of the STONE script and benchmarking data is available at https://github.com/TongZhou2017/STONE

## Ethics approval and consent to participate

Not applicable.

## Consent for publication

Not applicable.

## Competing interests

The authors declare that they have no competing interests.

## Notes

https://github.com/TongZhou2017/STONE

### Competing Interest Statement

The authors have declared no competing interest.

