## Supplemental Table 1 for "Integrating Mutation and Stop Signals for Improved RNA Structure Analysis and Insight Discovery"

### Table S1

| Title | Project ID | Dataset No. | Species No. | Method |
|---|---|---|---|---|
| In vivo genome-wide profiling of RNA secondary structure reveals novel regulatory features | PRJNA211324 | 4 | 1 | DMS Structure_seq |
| Genome-wide probing of RNA structure reveals active unfolding of mRNA structures in vivo | PRJNA196404 | 22 | 2 | DMS_seq |
| Mod-seq: high-throughput sequencing for chemical probing of RNA structure | PRJNA216133 | 8 | 1 | Mod-seq |
| Structural imprints in vivo decode RNA regulatory mechanisms | PRJNA257310 | 10 | 1 | icSHAPE |
| DMS-MaPseq for genome-wide or targeted RNA structure probing in vivo | PRJNA329551 | 22 | 3 | DMS-MaPseq |
| Keth-seq for transcriptome-wide RNA structure mapping | PRJNA503570 | 26 | 2 | Keth-seq |
| Comprehensive in vivo secondary structure of the SARS-CoV-2 genome reveals novel regulatory motifs and mechanisms | PRJNA645342 | 4 | 1 | SHAPE-MaP |
| A novel SHAPE reagent enables the analysis of RNAstructure in living cells with unprecedented accuracy | PRJNA646706 | 36 | 3 | SHAPE-Map |
| Genome-wide structure changes during neuronal differentiation drive gene regulatory networks | PRJNA658654 | 40 | 1 | icSHAPE |
|  | PRJNA946308 | 87 | 1 | scSPORT |
| RNA structure profiling at single-cell resolution reveals new determinants of cell identity | PRJNA946273 | 56 | 1 | scSPORT |
|  | PRJNA946372 | 26 | 1 | scSPORT |
| Predicting dynamic cellular protein–RNA interactions by deep learning using in vivo RNA structures | PRJNA608297 | 12 | 2 | icSHAPE |

### Table S1

| Dataset | Method | Small molecule | Enzyme | Strategy | Species | Quality Control | Reference Genome | Mapping Rate | No. of expressing genes | No. of genes with structural signals |
|---|---|---|---|---|---|---|---|---|---|---|
| FibroblastVitro_fastp_statistics | DMS_seq | DMS | Superscript III | Stop | Human | 98.81% | GCF_000001405.40, GRCh38.p14 | 82.13% | 51445 | 3938 |
| FibroblastVivo_fastp_statistics | DMS_seq | DMS | Superscript III | Stop | Human | 96.30% | GCF_000001405.40, GRCh38.p14 | 76.23% | 49415 | 2062 |
| k562vitro_fastp_statistics | DMS_seq | DMS | Superscript III | Stop | Human | 98.96% | GCF_000001405.40, GRCh38.p14 | 84.20% | 52229 | 4471 |
| K562vivo1_dms300_fastp_statistics | DMS_seq | DMS | Superscript III | Stop | Human | 97.88% | GCF_000001405.40, GRCh38.p14 | 82.02% | 53562 | 1673 |
| Vitro_30C_rep1_statistics | DMS_seq | DMS | Superscript III | Stop | Yeast | 98.49% | GCF_000146045.2, R64.sacCer3 | 76.14% | 6391 | 1558 |
| Vivo_rep2_statistics | DMS_seq | DMS | Superscript III | Stop | Yeast | 97.23% | GCF_000146045.2, R64.sacCer3 | 67.03% | 6376 | 1514 |
| 11_Scer_WT_Rep1_100mM_DMS | Mod-seq | DMS | Superscript II | Stop | Yeast | 75.81% | GCF_000146045.2, R64.sacCer4 | 89.93% | 4704 | 28 |
| v65_polyA_plus_icSHAPE_in_vitro_NAI_N3_Biological_Replicate_1_statistics | icSHAPE | NAI_N3 | Superscript III | Stop | Mouse | 89.41% | GCF_000001635.27, GRCm39 | 78.27% | 33211 | 5872 |
| HEK_293Ts_DMSvivo_genomeWide_fastp_statistics | DMS-MaPseq | DMS | TGIRT-III | mutation | Human | 99.57% | GCF_000001405.40, GRCh38.p14 | 78.60% | 50234 | 1988 |
| Scer_SSii_Mn_DMSvivo_Rep1_genomeWide_statistics | DMS-MaPseq | DMS | superscript II | mutation | Yeast | 99.49% | GCF_000146045.2, R64.sacCer3 | 81.29% | 6457 | 1871 |
| Scer_TGIRT_DMSvivo_Rep1_genomeWide_yeast_statistics | DMS-MaPseq | DMS | TGIRT-III | mutation | Yeast | 99.29% | GCF_000146045.2, R64.sacCer3 | 85.49% | 6444 | 1650 |
| D0_icShape_NAIN3_batch2_rep2 | icSHAPE | NAI-N3 | SSII MnCl2 | stop | Human | 51.77% | GCF_000001405.40, GRCh38.p14 | 91.30% | 42465 | 1946 |
| D7_icShape_NAIN3_batch2_rep2_statistics | icSHAPE | NAI-N3 | SSII MnCl2 | stop | Human | 59.21% | GCF_000001405.40, GRCh38.p14 | 92.73% | 43169 | 2674 |
| D8_icShape_NAIN3_batch2_rep2_statistics | icSHAPE | NAI-N3 | SSII MnCl2 | stop | Human | 56.72% | GCF_000001405.40, GRCh38.p14 | 91.54% | 44510 | 2624 |
| D14_icShape_NAIN3_batch2_rep2_statistics | icSHAPE | NAI-N3 | SSII MnCl2 | stop | Human | 52.48% | GCF_000001405.40, GRCh38.p14 | 91.08% | 44752 | 2662 |
| H9N_Homo_sapiens_statistics | icSHAPE | NAI-N3 | SuperScript III | stop | Human | 90.43% | GCF_000001405.40, GRCh38.p14 | 73.48% | 49403 | 4760 |
| HEK293N_Homo_sapiens_statistics | icSHAPE | NAI-N3 | SuperScript III | stop | Human | 96.85% | GCF_000001405.40, GRCh38.p14 | 94.54% | 50502 | 5468 |
| HeLaN_Homo_sapiens_statistics | icSHAPE | NAI-N3 | SuperScript III | stop | Human | 75.53% | GCF_000001405.40, GRCh38.p14 | 92.51% | 5711 | 1879 |
| HepG2N_Homo_sapiens_statistics | icSHAPE | NAI-N3 | SuperScript III | stop | Human | 97.17% | GCF_000001405.40, GRCh38.p14 | 96.12% | 49755 | 5354 |
| K562N_Homo_sapiens_statistics | icSHAPE | NAI-N3 | SuperScript III | stop | Human | 96.13% | GCF_000001405.40, GRCh38.p14 | 92.21% | 50173 | 4418 |
