## Supplemental Table 2 for "Integrating Mutation and Stop Signals for Improved RNA Structure Analysis and Insight Discovery"

### Table S2

| feature | description | importance rank in DMS model | importance rank in SHAPE model | unselect reason |
|---|---|---|---|---|
| rf_mutation_Count | mutation count number | 11 | 11 | |
| rf_mutation_Depth | sequencing depth | 8 | 5 | |
| rf_mutation_AC | The number of A-to-C base mutations | NA | NA | The incorporation of these features results in pronounced fluctuations in the Stone method's output during downsampling. As sequencing depth decreases, the AUC of the Stone method exhibits a marked decline, indicating that the inclusion of these features makes the Stone method more sensitive to variations in sequencing depth. |
| rf_mutation_AG | The number of A-to-G base mutations | NA | NA | |
| rf_mutation_AT | The number of A-to-T base mutations | NA | NA | |
| rf_mutation_CA | The number of C-to-A base mutations | NA | NA | |
| rf_mutation_CG | The number of C-to-G base mutations | NA | NA | |
| rf_mutation_CT | The number of C-to-T base mutations | NA | NA | |
| rf_mutation_GA | The number of G-to-A base mutations | NA | NA | |
| rf_mutation_GC | The number of G-to-C base mutations | NA | NA | |
| rf_mutation_GT | The number of G-to-T base mutations | NA | NA | |
| rf_mutation_TA | The number of T-to-A base mutations | NA | NA | |
| rf_mutation_TC | The number of T-to-C base mutations | NA | NA | |
| rf_mutation_TG | The number of T-to-G base mutations | NA | NA | |
| rf_mutation_ins | The number of insertion mutations | 15 | 15 | |
| rf_mutation_del | The number of deletion mutations | 12 | 7 | |
| pipe_truncation_ShapeScore | The SHAPE score of the Stop strategy | NA | NA | These values contribute to overfitting in the Stone model. |
| pipe_truncation_ChrPos | Position information | NA | NA | |
| pipe_truncation_RT | stop count number | 5 | 3 | |
| pipe_truncation_BD | sequencing depth | NA | NA | The values of pipe_truncation_BD and rf_mutation_Depth are identical, indicating that they represent the same concept. |
| base_A | The number of A bases | 9 | 2 | |
| base_T | The number of T bases | 13 | 9 | |
| base_G | The number of G bases | 14 | 6 | |
| base_C | The number of C bases | 10 | 10 | |
| rate_AT | The ratio of A-to-T base mutations:rf_mutation_AT/rf_mutation_Depth | NA | NA | The incorporation of these features results in pronounced fluctuations in the Stone method's output during downsampling. As sequencing depth decreases, the AUC of the Stone method exhibits a marked decline, indicating that the inclusion of these features makes the Stone method more sensitive to variations in sequencing depth. |
| rate_AG | The ratio of A-to-G base mutations:rf_mutation_AG/rf_mutation_Depth | NA | NA | |
| rate_AC | The ratio of A-to-C base mutations:rf_mutation_AC/rf_mutation_Depth | NA | NA | |
| rate_TA | The ratio of T-to-A base mutations:rf_mutation_TA/rf_mutation_Depth | NA | NA | |
| rate_TG | The ratio of T-to-G base mutations:rf_mutation_TG/rf_mutation_Depth | NA | NA | |
| rate_TC | The ratio of T-to-C base mutations:rf_mutation_TC/rf_mutation_Depth | NA | NA | |
| rate_GA | The ratio of G-to-A base mutations:rf_mutation_GA/rf_mutation_Depth | NA | NA | |
| rate_GT | The ratio of G-to-T base mutations:rf_mutation_GT/rf_mutation_Depth | NA | NA | |
| rate_GC | The ratio of G-to-C base mutations:rf_mutation_GC/rf_mutation_Depth | NA | NA | |
| rate_CA | The ratio of C-to-A base mutations:rf_mutation_CA/rf_mutation_Depth | NA | NA | |
| rate_CT | The ratio of C-to-T base mutations:rf_mutation_CT/rf_mutation_Depth | NA | NA | |
| rate_CG | The ratio of C-to-G base mutations:rf_mutation_CG/rf_mutation_Depth | NA | NA | |
| rate_A | The ratio of A base:base_A/rf_mutation_Depth | 2 | 4 | |
| rate_T | The ratio of T base:base_T/rf_mutation_Depth | 7 | 13 | |
| rate_C | The ratio of C base:base_C/rf_mutation_Depth | 1 | 14 | |
| rate_G | The ratio of G base:base_G/rf_mutation_Depth | 6 | 12 | |
| rate_mut | mutation ration:rf_mutation_Count/rf_mutation_Depth | 3 | 8 | |
| rate_stop | stop ration:pipe_truncation_RT/pipe_truncation_B | 4 | 1 | |
